# Travelling waves in somitogenesis: collective cellular properties emerge from time-delayed juxtacrine oscillation coupling

**DOI:** 10.1101/297671

**Authors:** Tomas Tomka, Dagmar Iber, Marcelo Boareto

## Abstract

The sculpturing of the vertebrate body plan into segments begins with the sequential formation of somites in the presomitic mesoderm (PSM). The rhythmicity of this process is controlled by travelling waves of gene expression. These kinetic waves emerge from coupled cellular oscillators and sweep across the PSM. In zebrafish, the oscillations are driven by autorepression of *her* genes and are synchronized via Notch signalling. Mathematical modelling has played an important role in explaining how collective properties emerge from the molecular interactions. Increasingly more quantitative experimental data permits the validation of those mathematical models, yet leads to increasingly more complex model formulations that hamper an intuitive understanding of the underlying mechanisms. Here, we review previous efforts, and design a mechanistic model of the *her1* oscillator, which represents the experimentally viable *her7;hes6* double mutant. This genetically simplified system is ideally suited to conceptually recapitulate oscillatory entrainment and travelling wave formation, and to highlight open questions. It shows that three key parameters, the autorepression delay, the juxtacrine coupling delay, and the coupling strength, are sufficient to understand the emergence of the collective period, the collective amplitude, and the synchronization of neighbouring Her1 oscillators. Moreover, two spatiotemporal time delay gradients, in the autorepression and in the juxtacrine signalling, are required to explain the collective oscillatory dynamics and synchrony of PSM cells. The highlighted developmental principles likely apply more generally to other developmental processes, including neurogenesis and angiogenesis.

## 1. Introduction

The body axis of vertebrates is segmented into anatomical modules consisting of vertebrae and their associated muscles. This segmentation has a key role in defining the mode of locomotion of an animal (Ward & Mehta, 2011). The embryonic precursors of the segments are called somites. The bilateral symmetric pairs of somites form in a process referred to as somitogenesis (Oates et al., 2012; Hubaud & Pourquié, 2014). At the same time as somitogenesis proceeds, the body axis is elongated posteriorly at the so-called tail bud by proliferation of presomitic mesoderm (PSM) cells originating from the marginal zone in zebrafish and frogs or the primitive streak in chicken and mice. Periodically, a pair of somites buds off from the anterior end of the PSM, which lies on both sides of the neural tube. In zebrafish, a new pair of somites is formed in the anterior trunk constantly every 23 min at 28.5℃ until this rhythm is gradually slowed down during the tail segmentation (Schröter et al., 2008). The rhythmicity is controlled by travelling waves of gene expression which sweep from the tail bud to the anterior end of the PSM (Palmeirim et al., 1997). The travelling waves emerge from the coordinated oscillation in individual cells (Jiang et al., 2000). In the following, we will first discuss a well-established theoretical model of somitogenesis, the clock-and-wavefront model. We will then in detail report what is known about the individual cellular oscillator, the coupling between neighbouring cells and the emergence of the travelling waves.

### 1.1 Rhythmical formation of somites

The most cited model for the sequential formation of somites is the clock-and-wavefront model (Cooke & Zeeman, 1976; Oates et al., 2012; Hubaud & Pourquié, 2014). It postulates that cells located at the anterior boundary of the PSM, the wavefront, undergo an abrupt change in cellular properties triggered by a periodic segmentation signal of the clock (Cooke & Zeeman, 1976). This clock, the first component of the model, has been shown to entail the oscillatory expression of genes involved in the Notch, WNT and FGF signalling pathways (Palmeirim et al., 1997; Dequeant et al., 2006). The most prominent examples are genes of the *Hes/her*family (Palmeirim et al., 1997; Cooke, 1998; Bessho et al., 2003). The oscillations in HES/Her are driven by time-delayed autorepression (Lewis, 2003; Monk, 2003). The corresponding time delay originates from mRNA splicing and nuclear export processes (Takashima et al., 2011; Hoyle & Ish-Horowicz, 2013). In principle, the oscillations in individual cells are synchronized perpendicularly to the body axis, such that they form travelling waves propagating in the anterior direction (Oates et al., 2012; Hubaud & Pourquié, 2014). Strictly, the bilateral symmetric travelling waves acquire a folded shape in zebrafish, termed a chevron, which is thought to arise from mechanical forces (Rost et al., 2014). The underlying synchronicity of neighbouring *Hes/her*oscillators is mediated by juxtracrine Notch signalling (Jiang et al., 2000; Lewis, 2003; Liao & Oates, 2016) and is discussed in section 1.2.

The second component of the clock-and-wavefront model, the wavefront, has been hypothesised to be defined via a threshold in the FGF or WNT gradients, which decline towards the anterior end of the PSM (Dubrulle et al., 2001; Sawada et al., 2001; Aulehla et al., 2003, 2008; Dunty et al., 2008; Naiche et al., 2011; Bajard et al., 2014). The FGF signalling gradient forms because the corresponding ligand-expressing gene *fgf8*is transcribed only in the tail bud and its mRNA is gradually decaying in the presomitic cells that are left behind by the proliferating tail bud (Dubrulle & Pourquié, 2004). More generally, this is referred to as a ‘gradient by inheritance’ - established by simultaneous cell flow and ligand mRNA or protein decay - and such a mechanism is thought to also govern the graded expression of *wnt3A*in the PSM (Aulehla & Pourquié, 2010; Bajard et al., 2014). Due to continuous axis elongation, the source of the ligand in the tail bud is progressively drawn further away from the anterior. The wavefront, presumably specified by a threshold in one of these gradients, is shifted accordingly. An opposing retinoic acid gradient shows highest concentration in the somites and is thought to antagonize the FGF gradient by mutual inhibition (Niederreither et al., 1997; Diez del Corral et al., 2003; Goldbeter et al., 2007; Jörg et al., 2016). Currently, this interpretation of the wavefront is largely debated: it has been proposed that the wavefront is implicitly defined by the clock, rather than being an independent entity (Lauschke et al., 2013).

According to the clock-and-wavefront model, segmentation is initiated when the clock signal reaches the moving wavefront (Cooke & Zeeman, 1976). This leads to a transition of the synchronized presomitic cell patches to distinct blocks of epithelial cells, the somites, that bud off from the PSM (Saga, 2012). The Mesp genes are essential in triggering this mesenchymal-to-epithelial transition (Saga et al., 1997; Takahashi et al., 2000; Sawada et al., 2000). In mice, *Mesp2* has been linked to both, the clock and wavefront, being positively regulated by Notch and negatively regulated by FGF signalling (Yasuhiko et al, 2006; Oginuma el al., 2008; Naiche et al., 2011). In zebrafish, expression of *mesp* genes seems to be similarly controlled by the wavefront, but the connection to the clock remains unclear (Sawada et al., 2001; Bajard et al., 2014; Wanglar et al., 2014; Yabe & Takada, 2016).

### 1.2. Architecture and synchrony of coupled Her oscillators

The autorepressive *Hes* and *her* genes lie at the core of the PSM oscillators in mice and zebrafish, respectively. These genes encode basic helix loop helix (bHLH) transcriptional repressors. In zebrafish, which serves us as a biological model organism, either of the two proteins Her1 and Her7 represses both corresponding genes in a redundant manner (Oates & Ho, 2002; Henry et al., 2007; Giudicelli et al., 2007). Another bHLH factor gene, *hes6*, is expressed in an FGF-dependent posterior-to-anterior gradient in the PSM (Kawamura et al., 2005). Her1, Her7 and Hes6 form various homo- and heterodimers, of which only Her1:Her1 and Her7:Hes6 are active repressors, targeting the promoters of *her1*, *her7* and *deltaC* (Schröter et al., 2012). The latter is the link that couples neighbouring oscillators: due to periodic repression by active Her dimers, *deltaC* is expressed cyclically and activates the *her* genes in neighbouring cells via Notch signalling (Jiang et al., 2000; Lewis, 2003; Liao & Oates, 2016). PSM cells have been shown to oscillate autonomously when uncoupled or isolated, although with a lower precision and persistence (Maroto et al., 2005; Masamizu et al., 2006; Webb et al., 2016).

The fact that the cells of the PSM synchronize their oscillations perpendicular to the direction of propagation of the travelling waves is intriguing. This synchronization requires cell-cell contact and is mediated by the coupling via Notch signalling (Jiang et al., 2000; Maroto et al., 2005; Horikawa et al., 2006; Riedel-Kruse et al., 2007; Özbudak & Lewis, 2008). However, it has been shown that the synchrony is initiated in the presumptive mesoderm ring independently of Notch signalling (Riedel-Kruse et al., 2007). In Notch pathway mutants, a few intact anterior somites are formed, but as segmentation proceeds to the posterior, severe defects occur (Jiang et al., 2000). The widely-supported desynchronization hypothesis has been founded on this observation and states that the essential role of Notch signalling is to maintain the initial synchrony (Jiang et al., 2000; Riedel-Kruse et al., 2007; Özbudak & Lewis, 2008; Liao & Oates, 2016).

Theoretical models show that coupling via Notch signalling can lead to either synchronization or salt-and-pepper pattern of oscillation phases, depending on the time delay involved in Notch signalling (Lewis, 2003; Tiedemann et al., 2007; Ay et al., 2013). Mathematical formulations show that the collective period of coupled oscillators might differ from the period of uncoupled oscillators, depending in a non-trivial way on both, the coupling strength and the coupling delay (Niebur et al., 1991; Morelli et al., 2009; Herrgen et al., 2010; Wang et al., 2014). In the PSM, the collective period is a local property and related measurements are discussed in section 1.3.

DeltaD, another ligand of the Notch pathway, is not expressed cyclically but in a decreasing gradient from posterior to anterior (Holley et al., 2002; Mara et al., 2007; Wright et al., 2011). In contrast to DeltaC, DeltaD is thought to cis-inhibit Notch (Matsuda & Chitnis, 2009). It has been suggested that DeltaD, unable to activate Notch by itself, could potentiate signalling via DeltaC by heterodimerization (Wright et al., 2011).

### 1.3 Travelling waves of gene expression

In zebrafish, the most prominent cyclically expressed genes that constitute the travelling wave are *her1*, *her7* and *deltaC* (Krol et al., 2011; Oates et al., 2012). From the posterior to the anterior, the travelling wave is slowing gradually while its wavelength decreases (Palmeirim et al., 1997; Giudicelli et al., 2007; Figure 1). Initially, the clock-and-wavefront model was thought to entail a genetic oscillator with a single, well-defined period that governs the rhythmicity of the segmentation clock (Cooke & Zeeman, 1976; Oates et al.; 2012). This simplistic picture has since been revisited with emphasis on the distinction between different notions of periods involved in the process.

**Figure 1.**
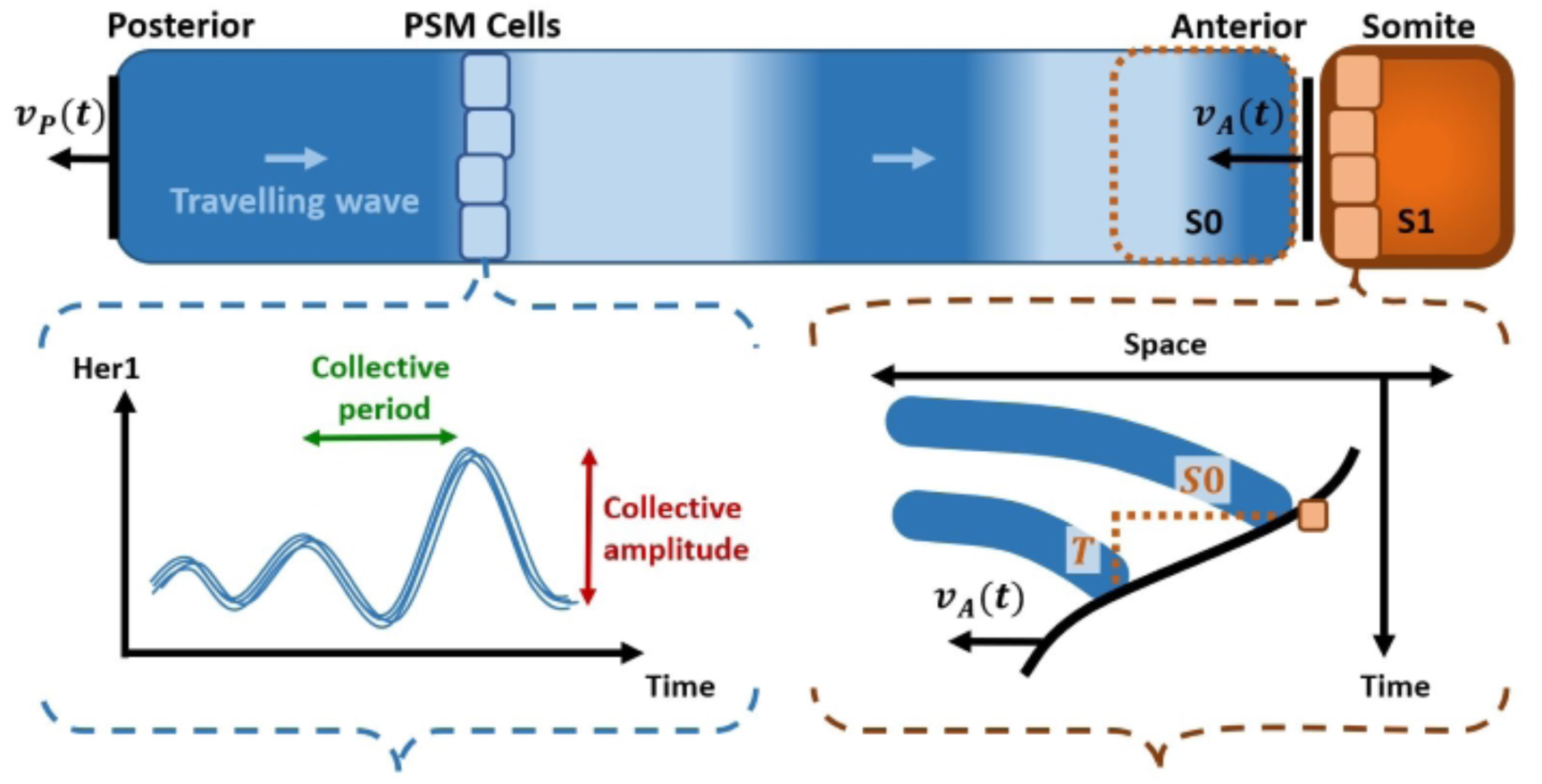
Schematic representation of somitogenesis from the reference frame of individual cells. In zebrafish, the anterior boundary of the PSM approaches the cells faster than the posterior boundary recedes due to proliferation; satisfying v_A_(t) > v_P_(t) during the entire process (Soroldoni et al., 2014). This leads to the shrinkage of the PSM over time (Soroldoni et al., 2014). Consequently, a Doppler effect is experienced at the anterior boundary when receiving the signal of the travelling wave (Soroldoni et al., 2014). The individual cellular oscillators will synchronize perpendicularly to the travelling wave direction, attaining a collective period and amplitude of cyclic components such as Her1 increase as the anterior boundary is approaching (bottom left; Delaune et al., 2012; Shih et al., 2015). The forming somite’s size S0, the somitogenesis period T and the velocity of the anterior boundary v_A_ are interdependent (bottom right; Jörg et al., 2015).

On a tissue level, one observes that the segmentation or somitogenesis period, the period with which the somites are formed, is equivalent to the period at which the travelling waves reach the wavefront (Soroldoni et al., 2014). This period is lengthened by the autorepression delay, as well as shortened in zebrafish or lengthened in mouse by Notch signalling (Harima et al., 2013; Kim et al., 2011; Herrgen et al., 2010; Webb et al., 2016; Liao et al., 2016). The segmentation is accelerated compared to the frequency at which travelling waves exit the tailbud due to a Doppler effect: relative to the tailbud, the anterior boundary is moving towards the approaching travelling waves and thus registers an increased frequency of wave signals (Soroldoni et al., 2014; Figure 1).

The size of the forming somite, referred to as *S*0, has been determined to be half of the wavelength at the PSM-somite border; the formation of *S*0 is completed when a full period of oscillation is cycled at this border (Shih et al., 2015; Figure 1). Given a local estimate for the velocity of the anterior PSM boundary 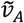, the somite size *S* is approximated as 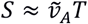where *T* is the somitogenesis period (Jörg et al., 2015; Figure 1 bottom right). There is also evidence suggesting that the somite size is linked to the phase gradient (Lauschke et al., 2013, Shih et al., 2015). However, the mechanism of oscillation arrest in *S*0, or the nature of the wavefront, remains elusive (Lauschke et al., 2013; Shih et al., 2015; Jörg et al., 2015). Furthermore, the segmentation scales with the body size (Cooke, 1975; Tam, 1981). Scaling of somite size, oscillation phase and travelling wave velocity has also been recently reported to occur during an *ex vivo* segmentation process, but the mechanism remains unknown (Lauschke et al., 2013).

### 1.4. Individual cell dynamics induce travelling wave formation

On the level of an individual cellular oscillator, single cell analyses reveal that both, the locally collective period and amplitude of Her1, are increasing as the cell is flowing across the PSM (Delaune et al., 2012; Shih et al., 2015; Figure 1). Several computational models show that in the presence of such a gradient in oscillation period, travelling waves emerge (Kærn et al., 2000; Jaeger & Goodwin, 2001; Tiedemann et al., 2007; Uriu et al., 2009; Ay et al., 2014). Mechanistically, a gradient in oscillation period, and therefore travelling waves, can be established by imposing a gradient of a model parameter, such as intronic delays or degradation rates (Tiedemann et al., 2007; Uriu et al., 2009; Ay et al., 2014). Hypothetically, such a reaction rate could be regulated by paracrine signalling, namely the FGF/WNT gradient present in the PSM. Because single-cell data is scarce and reaction rate estimates broad, these models still lag in accuracy behind non-mechanistic descriptions, which for instance can approximate somite size reasonably well (Morelli et al., 2009; Herrgen et al., 2010; Murray et al., 2013; Jörg et al., 2015; 2016).

In summary, the travelling waves observed in the PSM carry a repetitive segmentation signal, which is read out at the moving wavefront (Oates et al., 2012; Hubaud & Pourquié, 2014; Soroldoni et al., 2014; Shih et al., 2015). The formation of these travelling waves is based on three main principles: the oscillation of autorepressive *Hes/her* genes (Palmeirim et al., 1997; Cooke, 1998; Bessho et al., 2003; Lewis, 2003; Monk, 2003), the local synchronization of neighbouring *Hes/her* oscillators via Notch signalling (Jiang et al., 2000; Lewis, 2003; Giudicelli et al., 2007; Liao & Oates, 2016), and an increasing gradient of oscillation period, which defines the direction of propagation of the travelling wave (Kærn et al., 2000; Jaeger and Goodwin, 2001; Tiedemann et al., 2007; Uriu et al., 2009; Ay et al., 2014). Each of these principles has been studied in different mechanistic models of largely varying complexity (Lewis, 2003; Monk, 2003; Tiedemann et al., 2007; Uriu et al., 2009; Hester et al., 2011; Ay et al., 2014).

Here, we develop a parsimonious mechanistic model, corresponding to the *her7;hes6* double mutant, that can account for the three principles mentioned above. For that, we revisit a simple model of the zebrafish *her1* oscillator proposed by Julian Lewis (Lewis, 2003). Lewis’ representation combines both, the *her1* and the *her7* oscillator and couples neighbouring cells via *deltaC* (Lewis, 2003). In a similar fashion, we couple only the *her1* oscillators, similarly as previously done in analytical studies (Wang et al., 2014), but our model incorporates only two time delays, instead of three. Using this reductive approach, we can recapitulate the principles that govern the travelling wave. We find that both, the dynamics of the individual cellular oscillator and the synchronization of neighbouring PSM cells, are modulated by three key parameters of the model: the *Hes/her* autorepression delay, the intercellular coupling delay, and the coupling strength between neighbouring cells. These insights allow us to discuss the general role of Notch-mediated oscillation coupling in differentiation processes involved not only in somitogenesis, but also in other developmental processes such as neurogenesis and angiogenesis.

## 2. Methods

### 2.1. A parsimonious model of the zebrafish Her1 oscillator

The travelling waves occurring in the PSM are induced by the dynamics of individual cellular oscillators (Tiedemann et al., 2007; Uriu et al., 2009; Ay et al., 2014). The basis of a computational model of the travelling wave is therefore the cellular oscillator, which is driven by two negative feedback loops in zebrafish, one over the Her1 homodimer and one over the Her7:Hes6 heterodimer (Schröter et al., 2012). The two loops are thought to be redundant and the *her7;hes6* double mutant is segmented normally in most cases (Schröter et al., 2012). A reduced model, incorporating the autorepressive loop of *her1* only, should therefore be capable, in principle, to represent the PSM oscillator adequately. A mathematical representation of this reduced model of the uncoupled, or autonomous, zebrafish oscillator has been proposed as two delayed differential equations (Lewis 2003, Figure 2):

**Figure 2:**
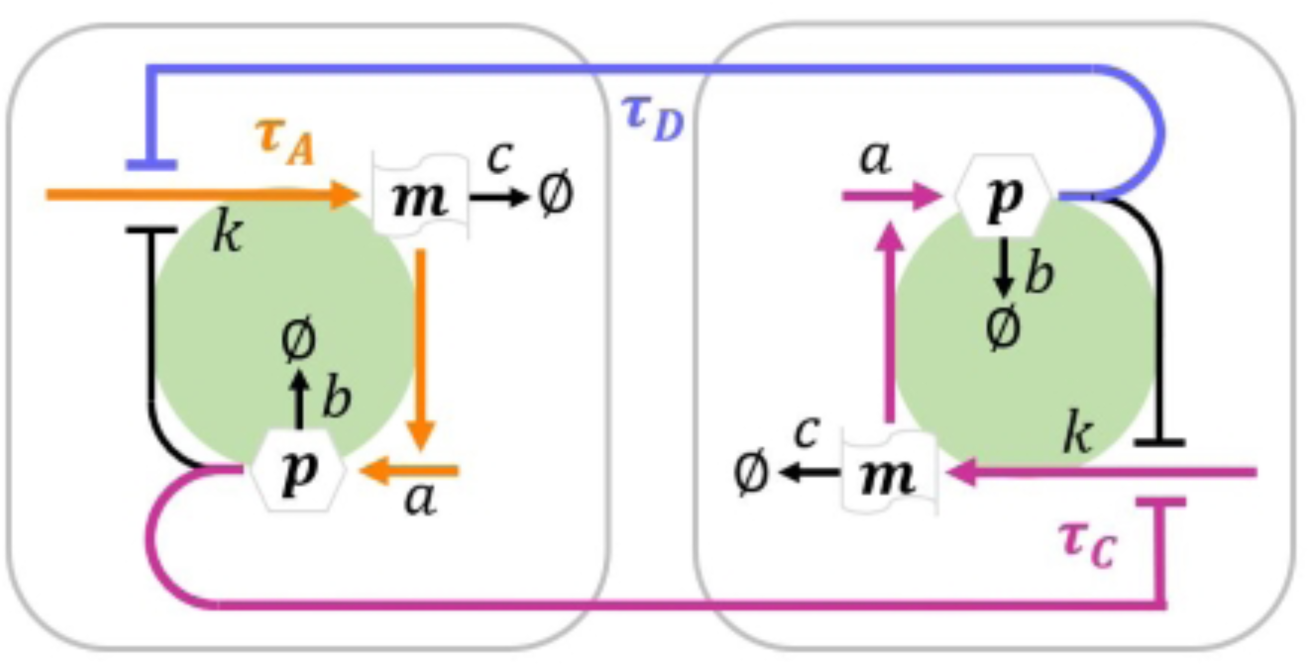
Graphical representation the model for the cellular oscillators in the PSM. Each cell contains a loop over the autorepressive protein, p, and its mRNA, m,with an autorepression delay τ_A_, which delays mRNA transcription. The oscillators are coupled between neighbours by mutual repression with a coupling delay τ_C_; it is the sum of the autorepression delay τ_A_and the deltaC expression delay τ_D_(Eq. 5). This coupling delay represents the time needed to transmit the phase information between cyclic protein levels in neighbouring cells. The kinetic rates a, b, c, and K, represent protein translation, protein degradation, mRNA degradation, and mRNA transcription, respectively. The model is mathematically described by Eq. 3,4.

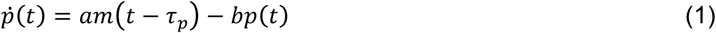

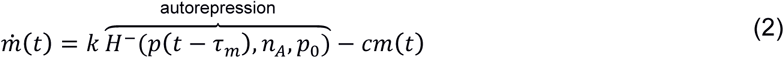

The two components of the model are the cytosolic mRNA, *m*, and the nuclear protein, *p*, of Her1. The nuclear protein is produced at a translation rate *a* = 4.5 min^−1^and degraded at a rate *b* = 0.23 min^−1^(Lewis, 2003). The transcription rate is given by *k* = 33 min^−1^and the mRNA is degraded at *c* = 0.23 min^−1^, the same degradation rate as for the protein (Lewis, 2003). A negative Hill function *H* ^−1^is used to model autorepression, with a Hill constant *p*_0_= 40 and a Hill coefficient *n*_A_= *2*. The translational and transcriptional delays have been estimated to be *τ*_p_≈ 2.8 min and τ_m_≈ 10 min, respectively (Lewis, 2003; Hanisch et al., 2013).

Additionally, we couple neighbouring oscillators via mutual repression, representing Notch signalling (Eq. 3, 4; Figure 2).

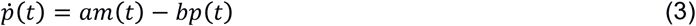

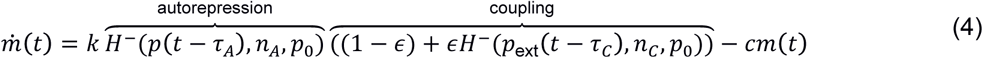

Different coupling mechanisms have been introduced, including very similar formulations (Lewis, 2003; Tiedemann et al., 2007; Kim et al., 2010; Wang et al., 2014). Lewis himself coupled neighbouring oscillators consisting of both, a *her1* and a *her7* circuit, and represented *deltaC* explicitly (Lewis, 2003). The transcription is repressed by the Notch signal, which depends in our model on the average Her1 concentration in neighbouring cells *p*_ext_. Again, we use a negative Hill function to model coupling via mutual repression, with a Hill constant *p*_0_= 40 and a Hill coefficient *n*_c_= 1. We introduced the parameter ϵ ∈ [0,1] to regulate coupling strength; as a default, we use maximal coupling, ϵ = 1. We also summarize the *her1* -related delays into one autorepression delay τ_A_, rendering the model more abstract, but simpler to understand. We considered τ_A_= 10 min as a default value. The expression of *deltaC* is delayed by translational and transcriptional delays, which have been estimated to be roughly 30 min and 10 min, respectively, resulting in a total *deltaC* expression delay τ_D_≈ 40 min (Giudicelli et al., 2007; Hanisch et al., 2013). These time delay estimates have been deduced indirectly from analyses of the spatial patterns of travelling waves and therefore should be used with care. The juxtacrine signalling delay via Notch, referred to as the coupling delay, is defined as

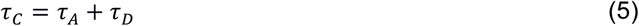

Consequently, its approximate duration is τ_c_≈ 50 min.

The zebrafish segmentation process is sensitive to body size and temperature (Cooke, 1975; Tam, 1981; Schröter et al., 2008). Additionally, the somitogenesis period of the *her7;hes6* double mutant is slower relative to the wild type (Schröter et al., 2012). Therefore, the reductive approach of our model allows us to make conceptual, but not quantitative, conclusions.

### 2.2. Cell-based simulations

To approximate the PSM locally, we use 4×4 cell patches on a quadratic lattice with periodic boundary conditions and homogeneous parameter values, similarly to previous studies (Tiedemann et al., 2007; Ay et al., 2013). We calculate the oscillation properties after the cells have reached a stable oscillating state. To measure synchronization of Her1 protein levels across such a tissue, we take the average pairwise Pearson’s *r* coefficient of Her1 concentration trajectories between neighbouring cells. The corresponding value lies between −1, for perfect anti-synchronization, and 1, for perfect synchronization. A white Gaussian noise is implemented proportional to the maximum level of the variable (protein or mRNA) times a control parameter η, such that η = 0 represents no noise and η = 1 represents a noise proportional to the maximum level observed.

To simulate travelling waves in the PSM, we use a domain of 4×50 cells on a quadratic lattice. We chose boundary conditions that correspond to the geometry of a cylinder and we induce parameter value gradients across the cylinder height spanning 50 cells, which represents the anterior-posterior axis. For simulations on dynamic domains, we adopt the approach of Ay et al. (2014): Firstly, we simulated 4×20 cells for 84 min. This domain represents the tail bud, which we assume to be homogeneous in parameter values, with a common period (Oates et al., 2012). Secondly, we divide the 4 posterior-most cells every 6 min for 150 min, until the entire 4×50 cell domain is filled (Ay et al., 2014). After a cell exits the tail bud the time delay parameters increase every 6 min as it moves toward the anterior boundary (as suggested by the results described below). Thirdly, we model stationary cell flow, by additionally removing the 4 anterior-most cells every 6 min (Ay et al., 2014). This stationary description is a simplification and does not allow for a quantitative description of the travelling wave (Soroldoni et al., 2014; Jörg et al., 2015).

### 2.3 Code availability

The delay equations (Eq. 3, 4) are computationally solved using Pydelay version 0.1.1 (Flunkert & Schoell, 2009). All source codes used in our simulations are presented as Jupyter notebooks (http://jupyter.org/) for easy visualization and are freely available on GitHub: https://github.com/ttomka/her1_somitogenesis

## 3. Results

### 3.1. Increasing the autorepression delay lengthens the period of autonomous

The Hes/her *oscillations.* The *Hes* and *her* genes drive the PSM oscillator in mice and zebrafish, respectively. Numerical and analytical results show that the period of the autonomous *Hes/her* oscillator depends on the autorepression delay and the protein and mRNA half-lives (Supplementary figure 1; Lewis, 2003; Monk, 2003; Hori et al., 2013). In our model, when the autorepression delay of *her1*, τ_A_, is lengthened, both, the autonomous period and amplitude increase (Figure 3A). There is a lower limit τ_A_≈ 7.5 min for the existence of oscillations, which depends on the kinetic rates and the Hill coefficient (Figure 3A; Hori et al., 2013; Novak & Tyson, 2008).

The relationship of the period and the delay can be understood intuitively: the autorepression peak must reside in the region where the oscillation amplitude is decreasing (Figure 3B). Therefore, the period must be at least twice as long as the autorepression delay. Lewis described this phenomenon mathematically: in the extreme case of a discrete on/off oscillation the autonomous period is given by *T*_a_= *2* τ_A_(Lewis, 2003). In reality, the mRNA and protein lifetimes also contribute to this period length (Lewis, 2003).

**Figure 3:**
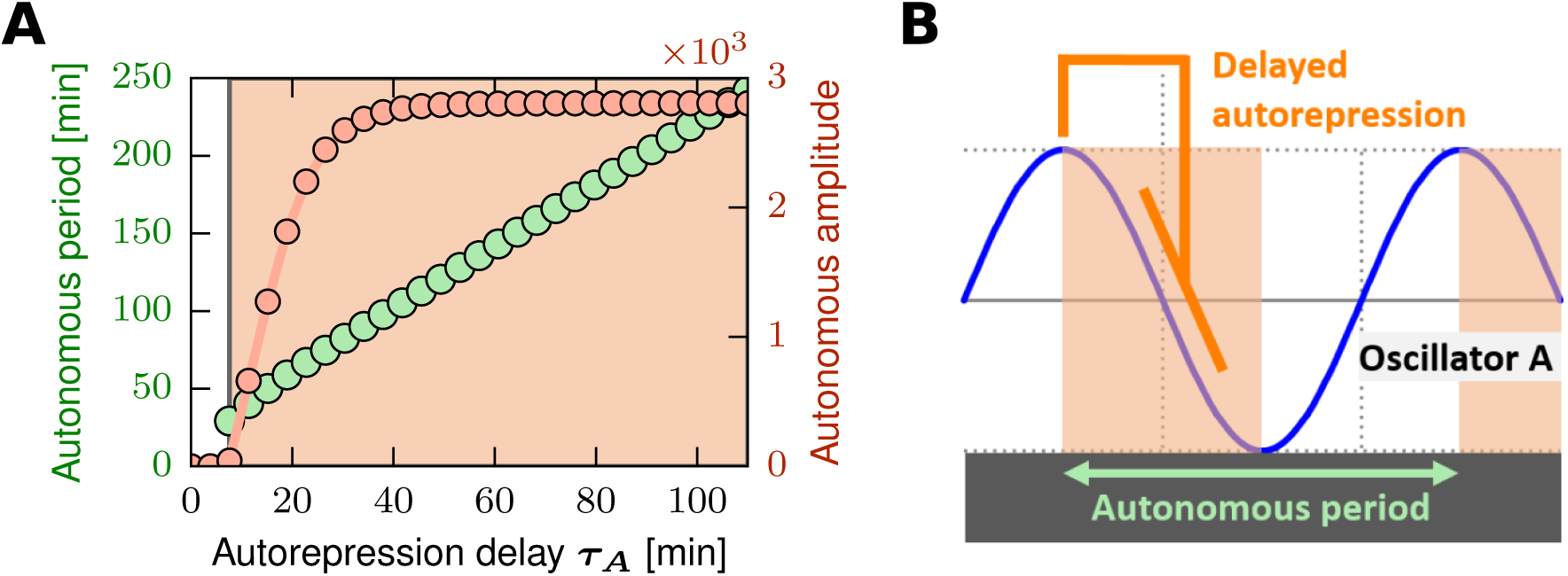
The autorepression delay can modulate the period and amplitude in an autonomous Her1 oscillator. (A) The period and amplitude of autonomous oscillations depend on the autorepression delay. (B) Graphical representation of the relationship of the autorepression delay and the autonomous period of the Her1 oscillator. An increase of the autorepression delay expands the orange region, where oscillation is sustained (see SI for details), because the maximal repression signal takes longer to shut-down her1 expression, which leads to a longer oscillation period.

### 3.2. Juxtacrine coupling controls cell-cell synchronization and robustness against noise

Riedel-Kruse et al. proposed that an initial synchrony of the cellular *her* oscillators is established in the presumptive mesoderm by simultaneous gene induction (Riedel-Kruse et al., 2007). In Notch pathway mutants, where oscillations are thought be autonomous, this synchrony is gradually lost (Jiang et al., 2000; Riedel-Kruse et al., 2007; Özbudak & Lewis, 2008; Liao & Oates, 2016). In our model, by setting all initial concentrations to zero and starting expression in all cells simultaneously, we observe a first peak of Her1 oscillation that is massively increased relative to the consecutive peaks, because the system initially has no memory of autorepression (Figure 4A). As time goes, in the absence of intercellular coupling, the autonomously oscillating cells lose their synchrony due to noise (Figure 4A).

**Figure 4:**
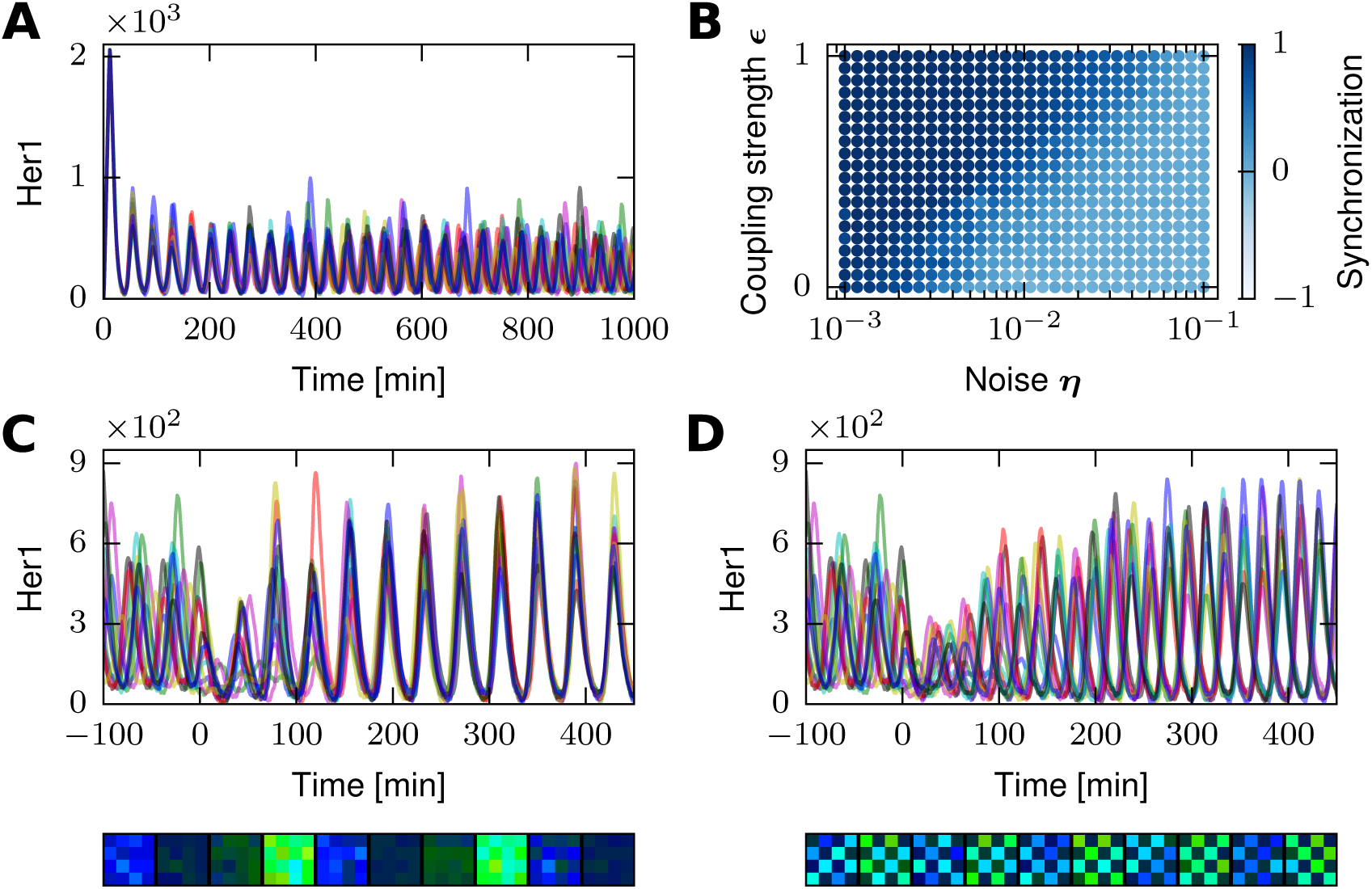
Coupling leads to robust cell-cell synchronization or dynamic salt-and-pepper patterning. (A) Simultaneous onset of Her1 expression leads to synchrony, which is lost over time in the absence of coupling. (B) Stronger coupling renders synchronization more robust, i.e., less sensitive to noise (average over 5 replicates). (C, D) At the endpoint of (A) which is now referred to as timepoint 0, the cells are coupled with coupling delays (C) τ_C_= 50 min (default value) or (D) τ_C_= 30min. The spatial patterns are displayed (bottom) for the last 90 minutes of the corresponding concentration trajectories (top). The noise level in (C, D) corresponds to the value of η = 10^−2^in (B); see Methods for details. All data has been recorded for 4×4 cell patches.

In the wild type, autonomous oscillators are coupled via Notch signalling, which increases robustness of synchronization (Jiang et al., 2000; Riedel-Kruse et al., 2007; Özbudak & Lewis, 2008; Soza-Ried et al., 2014; Jenkins et al., 2015; Liao & Oates, 2016). To investigate the effect of cell-cell coupling, we simulated a tissue of cells with different levels of noise and coupling strength, and we confirm that stronger coupling increases the robustness of synchronization, i.e. it renders the synchronization less sensitive to noise (Figure 4B). We further simulate how sudden coupling entrains oscillators which lost their synchrony in the presence of noise (Figure 4C and 4D). Consistent with previous studies (Lewis, 2003; Tiedemann et al., 2007; Morelli et al., 2009; Herrgen et al., 2010), we observe that the cells either synchronize (Figure 4C) or anti-synchronize (Figure 4D), depending on the value of the coupling delay. On a tissue level, the latter is also referred to as dynamical salt-and-pepper patterning and has been proposed to regulate the maintenance of neural progenitors during brain development (Shimojo et al., 2008; Kageyama et al., 2008).

Taken together, we observe in our model that the strength of juxtacrine coupling is regulating the robustness of cell-cell entrainment, while the delay of this coupling is regulating whether expression of *her1* in neighbouring cells is synchronized or anti-synchronized. In the following section, we examine how exactly the coupling delay influences the collective behaviour of *her1* oscillators.

### 3.3. Collective oscillatory dynamics critically depend on the coupling delay

We next evaluate the behaviour of our model for a range of coupling delays τ_c_. We observe alternating regions of synchronization and anti-synchronization (Figure 5A and 5B; for exemplary simulations see Figure 4C and 4D). In each region, the collective period increases monotonically (Figure 5A) and the collective amplitude describes a parabola with a local maximum (Figure 5B). A similar pattern has been reported partially in the context of the zebrafish PSM oscillator, *Hes/her* oscillations in neural differentiation, and a variety of synchronization phenomena across the natural sciences (Wang et al. 2014; Morelli et al., 2009; Herrgen et al., 2010; Momiji & Monk, 2009; Sadeghi & Valizadeh, 2014; Pavlides et al., 2015; Vanag et al., 2016; Wetzel et al., 2017). Nevertheless, in contrast to other mathematical descriptions, we do not observe oscillation death, a stable non-oscillating state, which has been hypothesized to occur between the synchronization and anti-synchronization phase space regions (Figure 5B, Shimojo et al., 2016). Furthermore, we observe in our *her7;hes6* double mutant model that for most coupling delays the collective period is higher than the autonomous period (Figure 5A), contrasting with measurements in wild type (Herrgen et al., 2010; Webb et al., 2016).

**Figure 5:**
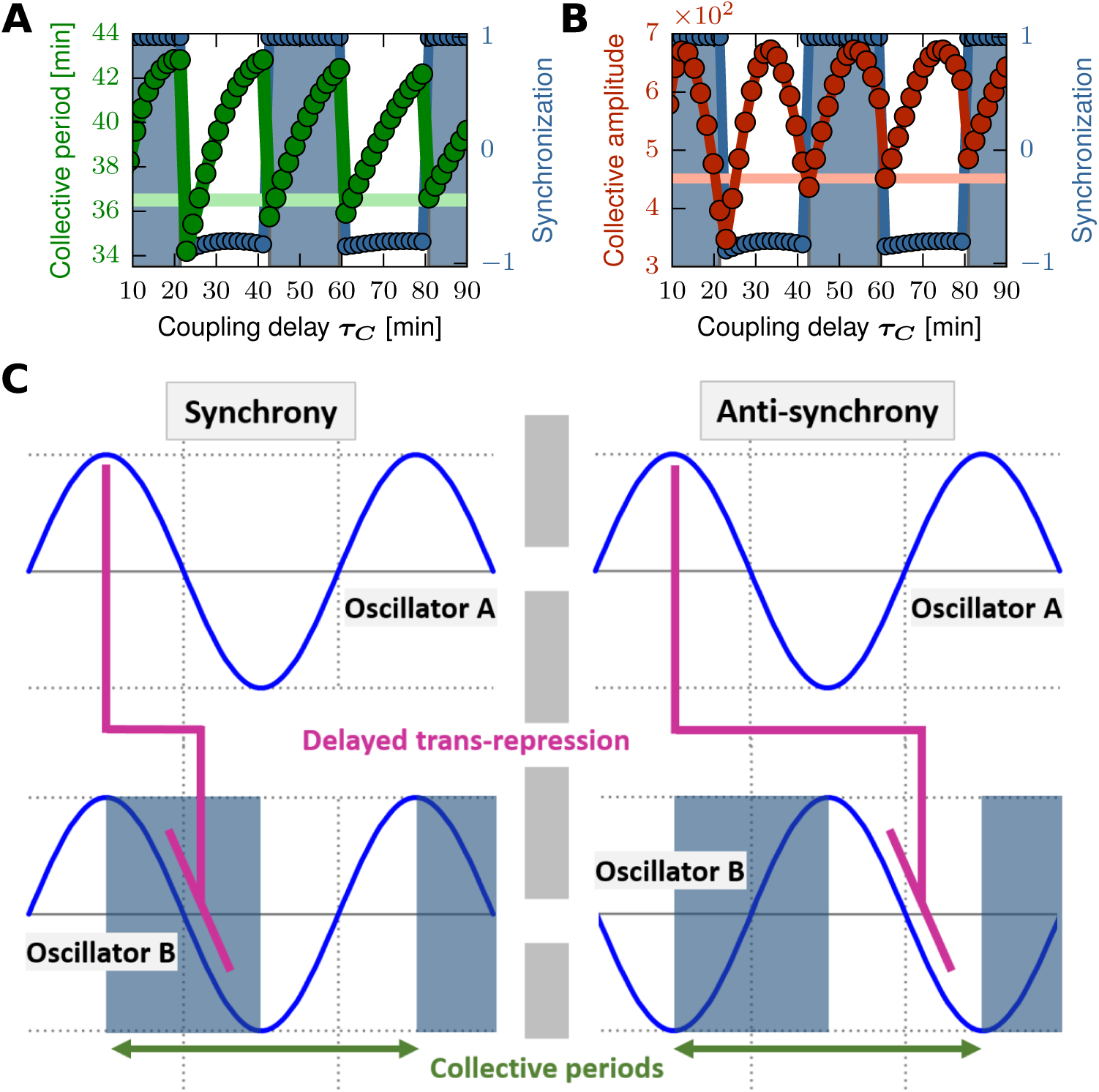
Collective period, collective amplitude and synchronization of Her1 oscillators critically depend on the coupling delay. (A, B) Dark blue points represent the synchronization (correlation in expression) between cells calculated by our model, while the blue region represents synchronized behaviour predicted by our heuristic argument (see SI for details). (A) Dark green points represent the collective period and (B) dark red points represent the collective amplitude of Her1 for different coupling delays. As a reference, we plot the autonomous period/amplitude of Her1 (light green/red line); data points correspond to 4×4 cell patches starting in a desynchronized initial condition (Figure 4A). (C) Graphical representation summarizes the dependence of synchronization and anti-synchronization on the collective period and the coupling delay. The peak of repression lies in the phase of the receiving oscillator where it decreases in amplitude; this determines the synchronization mode if the coupling delay and the collective period are given.

What is it that defines whether cells synchronize or not? Using the words of Lewis, *“the ‘push’ that the one cell delivers to its neighbour must be received in the correct part of the neighbour’s own oscillation cycle if it is to promote synchronized oscillation” (* Lewis 2003). We describe this “correct part” heuristically, as the time intervals where the oscillation amplitude is decreasing, given the neighbouring oscillators are aligned in-phase (blue regions in Figure 5; see SI for details). When the coupling delay does not meet this requirement, the phases will adjust to anti-synchrony, such that the “push” still occurs in the region where gene expression descends (Figure 5C, right). We notice that the crudely estimated coupling delay τ_c_≈ 50 min lies in the second region of synchronization (Figure 5A and 5B). In other words, the information is transmitted to the neighbouring cell with a delay bigger than one oscillation cycle.

In summary, when varying the coupling delay in our model, we observe alternating regions of synchronization and anti-synchronization with a repetitive pattern of collective periods and amplitudes (Figure 5).

### 3.4. Time delays control the collective behaviour of cellular oscillators across the PSM

We have shown above that for a given autorepression delay, modulations of the coupling will critically define collective properties of cellular *her1* oscillators (Figure 5). We have observed alternating regions of synchronization and anti-synchronization with respect to the coupling delay (Figure 5). Evaluating the combined effect of both, the autorepression and the coupling delay, we observe that the regions of synchronization and anti-synchronization gradually shift towards higher coupling delay values when increasing the autorepression delay (Figure 6A). Because the autorepression delay τ_A_is included in the coupling delay τ_c_(Eq. 5), τ_A_> τ_c_is infeasible (Figure 6A) and possible trajectories in the time delay phase space are restricted: the change in coupling delay must be greater than or equal to the change in the autorepression delay, Δτ_c_≥ Δτ_A_.

**Figure 6:**
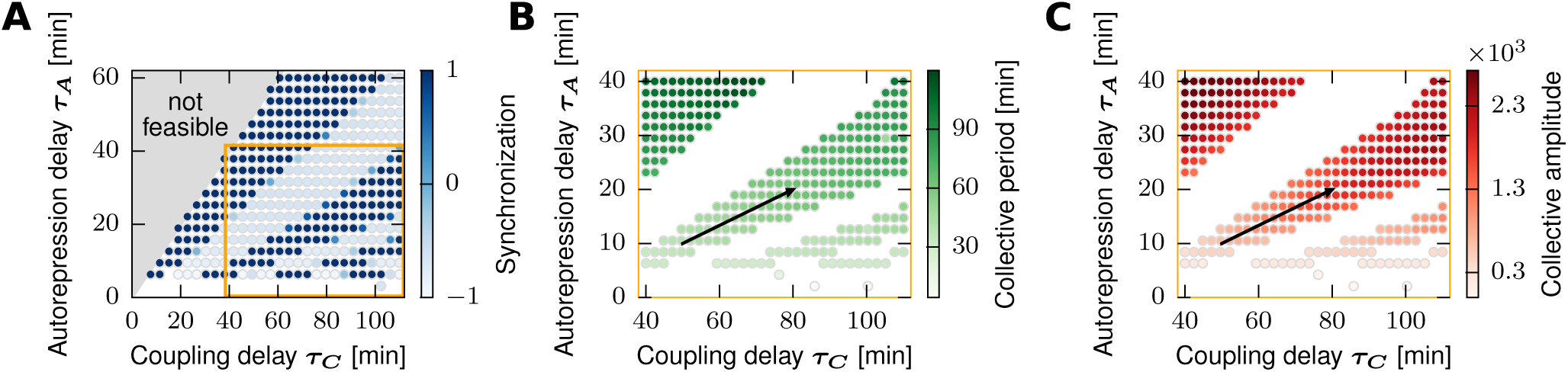
The time delay phase space of coupled Her1 oscillators reveals a path for synchronized oscillations with increasing collective periods and amplitudes. (A) Regions of synchronization (dark blue) and anti-synchronization (light blue) observed in the time delay phase space; the coupling delay contains the autorepression delay, which constrains the phase space (infeasible region coloured in light grey). (B, C) A subspace of (A) is shown (indicated by the orange frames); the anti-synchronized states are hidden (white region); the black arrows represent an exemplary path for cellular oscillator as it moves across the PSM (Figure 1), with increasing collective period (B) and amplitude (C) as indicated from experiments (Shih et al., 2015). All data points are averages corresponding to 4×4 cell patches starting in different desynchronized initial conditions (n=5).

Recently, experiments tracking the expression of single cells have revealed that the period of Her1 oscillations increases ~1.5-fold from the posterior to the anterior, while the amplitude roughly doubles (Shih et al., 2015). Concordantly with a previous mechanistic model (Ay et al., 2014), we observe that modulated time delays could achieve such variations across the PSM (Figure 6B and 6C). Moreover, to remain within the second region of synchronization (Figure 5A and 5B), both involved delays must increase as the oscillator travels across the PSM (Figure 6B and 6C). More precisely, according to our model, it is not sufficient to increase the coupling delay τ_c_only by lengthening the autorepression delay τ_A_, but an additional DeltaC-dependent time delay change Δτ_D_is needed, such that Δτ_C_= Δτ_A_+ Δτ_D_> Δτ_A_. For instance, the delays would change from τ_A,post_= 10 min and τ_c,post_= 50 min in the posterior to τ_A,ant_= 20 min and τ_c,ant_= 80 min in the anterior PSM (Figure 6B and 6C, black arrow); these values are taken as default in the following.

Importantly, we find that the other parameters of the *her1* oscillator are not suitable modulators of the collective period. Firstly, the collective period is not sensitive enough with respect to the kinetic rates of synthesis, translation and transcription (Supplementary figure 2; Ay et al., 2014; Wang et al., 2015). The same is true for the coupling strength, which furthermore affects robustness of synchronization (Supplementary figure 3, Figure 4A). Lastly, changes in degradation rates of Her1 protein or mRNA cannot increase both, the collective period and amplitude, simultaneously (Supplementary figure 2).

Taken together, our model suggests that gradients in the autorepression and coupling delays are critical to increase the collective period and amplitude as experimentally observed (Shih et al., 2015), while allowing neighbouring cells to remain in a synchronized condition.

### 3.5. Different mechanisms can lead to travelling waves in the presence of spatiotemporal time delay gradients

In Notch pathway mutants, the coupling between neighbouring cells is interrupted and the expression of *Hes/her* oscillates autonomously. In this case, their oscillatory dynamics that can be regulated on a large scale only by the autorepression delay (Figure 3, Supplementary figure S4). In contrast, in the presence of coupling, HES/Her oscillatory dynamics arise from a different mechanism (Figure 5): cells attain a locally collective period and amplitude, both of which can be modulated on a large scale only by changing both time delays of the system at the same time (Figure 6, Supplementary figure S4). In the following, we show that both mechanisms, coupled and autonomous oscillation, lead to travelling waves, in principle.

First, we simulate the temporal behaviour of a local group of homogeneous cells while increasing the time delays as suggested above (Figure 6A and 6B, black arrow), which leads to successively longer oscillations with higher amplitude in both scenarios, with autonomous and coupled oscillators (Figure 7A and 7B). When assuming continuous cell flow, the time a group of cells has spent travelling across the domain is also reflected by their position. It is unknown, whether the dynamics of the cellular oscillators are controlled by positional or temporal information (Oates et al., 2012; Shih et al., 2015). Both interpretations are possible.

**Figure 7:**
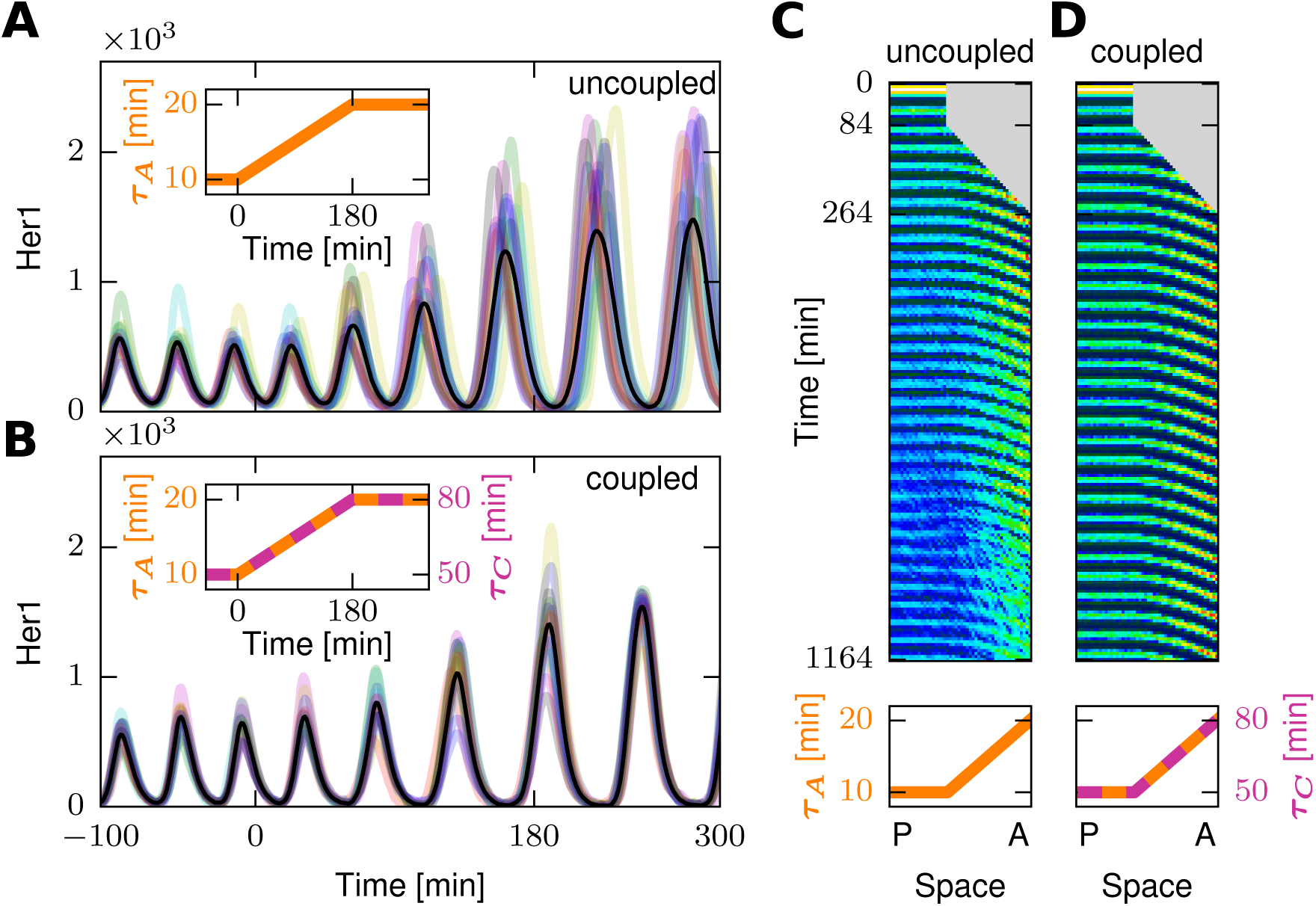
Travelling waves of autonomous or coupled oscillators. (A, B) Trajectories of autonomous (A) and coupled Her1 oscillations (B) in a group of 4×4 cells, in the presence of a temporal linear gradient in the autorepression delay τ_A_and the coupling delay τ_C_between the timepoints 0 min and 180 min (small embedded plots); black lines correspond to the average trajectories. (C, D) Kymographs of travelling wave series formed by autonomous (C) and coupled Her1 oscillators (D) under the influence of time delay gradients (bottom); at every time point ( 6 min intervals), the 4×50 tissue representing the PSM has been averaged over the medio-lateral axis; after 264 min, a stationary cell flow is maintained (for details refer to the Methods section). All time delay gradients correspond to the black arrow in Figure 6B and 6C. The noise level in the simulations corresponds to the value η = 10^−2^in Figure 4B (see Methods).

We observe that in the case of uncoupled oscillators, initial synchrony is gradually lost (Figure 7A), leading to severe irregularities in the wave pattern (Figure 7C). This corresponds well with experimentally observed posterior segmentation defects in Notch pathway mutants (Jiang et al., 2000). Our model suggests that in the case of autonomous cells, representing the Notch mutant, only a gradient in the autorepression delay τ_A_is needed to control the oscillatory dynamics (Figure 7A) and to form travelling waves. However, for coupled cells, i.e. when Notch signalling is present, changing the autorepression delay, which is also included in the coupling delay τ_c_, is not sufficient. Such a case would lead to a switch to a salt-and-pepper patterning regime (Supplementary figure S5). As we have hypothesized above, an additional DeltaC-dependent time delay change Δτ_D_is needed for the emergence of travelling waves (Figure 7D), such that Δτ_c_= Δτ_A_+ Δτ_D_> Δτ_A_.

Taken together, our analysis suggests that in the absence of cell-cell coupling, a delay in the *her* autorepression alone is sufficient to lead to travelling waves of *her* expression. In contrast, when the cells are coupled, an additional DeltaC-dependent delay is required in order to maintain the cells synchronized.

## 4. Discussion

### 4.1. A parsimonious mechanistic model recapitulates the travelling wave formation in the presomitic mesoderm

Recent cell-based models of somitogenesis have aimed to integrate explicitly the increasing amount of new experimental observations in a mechanistic network (Hester et al., 2011; Tiedemann et al., 2012; Ay et al., 2013). Incidentally, this resulted in increasingly complex models, compared to the first simple autorepression models of the segmentation clock (Lewis, 2003; Monk, 2003). We are convinced that examining minimal models can be an instructive approach to gain a more fundamental understanding of complex processes (Goldenfeld & Kadanoff, 1999). Therefore, we propose a simple model in a reduced form, compared to the model of Lewis (Lewis 2003; Eq. 1, 2). Our model includes only one autorepression cycle over *her1* and couples neighbouring oscillators via mutual repression (Eq. 3, 4). A similar model has been used for analytical purposes (Wang et al., 2014), but we incorporate only two time delays, instead of three. Our model corresponds to the *her7;hes6* double mutant. Because this mutant is segmented normally in most cases (Schröter et al., 2012), we assume that *her1* is sufficient to drive the cellular oscillators in the PSM. In appreciation of Julian Lewis, we show that his simple initial *her1* model is still remarkably powerful (Lewis, 2003; Lander, 2014). It is the key, which we use to understand the principles governing the travelling wave that controls the rhythmicity of the vertebrate segmentation. On this foundation, we offer a parsimonious mechanistic model, which recapitulates the properties of travelling waves in the PSM studied over decades in experiments and a variety of theoretical frameworks.

We show that the travelling wave formation in zebrafish can be understood by the control of three parameters: the autorepression delay, the coupling delay, and the coupling strength. These three parameters critically define three emergent local properties: the collective period, the collective amplitude, and the synchronization of neighbouring presomitic cells. As found in a more complex framework (Ay et al., 2014), we observe that a spatiotemporal gradient in the time delays can explain the cellular oscillator dynamics monitored *in vivo* (Figure 6B and 6C; Shih et al., 2015). Concomitantly, a synchronizing condition is maintained (Figure 6A).

### 4.2. Time delays in autonomous and coupled Her1 oscillators and their spatiotemporal control

We find that the oscillatory dynamics within the travelling wave in the PSM are best understood by examining the trajectory of the PSM cells through their time delay phase space (Figure 6). Firstly, we emphasize the role of the autorepression delay in modulating the period of the autonomous Her1 oscillator (Figure 3). A successive increase of this delay in autonomous oscillators will lead to a period and amplitude profile, which is on the same scale as measured experimentally in the wild type (Figure 7A and Supplementary figure S4; Shih et al., 2015). An effective translational delay gradient of Her1 has been measured experimentally (Ay et al., 2014). The desynchronization hypothesis states that uncoupled oscillators will lose their synchrony successively; this synchrony is established earlier in development by simultaneous gene induction of *her1* (Figure 4A, 7A and 7C; Jiang et al., 2000; Riedel-Kruse et al., 2007; Özbudak & Lewis, 2008; Liao & Oates, 2016).

The second time delay in our model, the coupling delay, represents the time needed for coupled oscillators to exchange Her1 oscillation phase information. This delay is not relevant in Notch pathway mutants, where juxtacrine coupling is interrupted and cells oscillate autonomously. We conclude that due to the long coupling delay of roughly 50 min, the information exchange in the *her7;hes6* double mutant takes longer than one oscillation cycle (second region of synchronization in Figure 5 and 6A). The experimental measurement of this coupling delay is inherently difficult. But for any value between 43 min and 60 min, the system remains in the same region of synchronization (Figure 5), which will lead to the same qualitative conclusions. In general, the coupling delay is a potent regulator of collective period and amplitude, and defines sharp transitions between synchronization and salt-and-pepper patterning (Figure 4C, 4D and 5; see further discussion below). Based on our numerical simulations (Figure 5A, 7C and 7D), we expect that in the *her7;hes6* double mutant the autonomous oscillators are faster than the coupled oscillators. While the same is true for mice (Kim et al., 2011), this contrasts with the zebrafish wild type, where the autonomous oscillators are slower than the coupled oscillators (Herrgen et al., 2010; Webb et al, 2016; Ay et al., 2013). It is a limitation of our model that it cannot approximate the wild type in this respect and it remains an open question what modulates these relative period differences mechanistically.

We notice an intuitive similarity in the behaviour of coupling and autorepression feedback loops: to sustain oscillation, the peak of inhibition must occur in the time intervals where the amplitude of the receiving oscillator is decreasing. Either the receiving oscillator is the sending oscillator itself (Figure 3B) or its neighbour (Figure 5C).

Because the coupling delay incorporates the autorepression delay, changes in coupling delay must be greater or equal to the changes in autorepression delay for an individual cell that moves through the time delay phase space (Figure 6A). We have discussed that for autonomous Her1 oscillators it is only possible to modulate the autorepression delay to achieve reasonable period and amplitude gradients (Figure 7A and Supplementary figure S4). This mechanism, however, is not applicable for the coupled Her1 oscillators (Supplementary figure S5), which modulate their autonomous period additionally by exchanging phase information with their neighbours. For the coupled oscillators, our model indicates only one mechanism to achieve gradients in collective period and amplitude, and to remain in the same region of synchronization (Figure 6): besides the autorepression delay gradient, an additional spatiotemporal gradient in the *deltaC* expression delay is required to form travelling waves (Figure 7D). These predictions are in line with a more complex model proposed by others (Ay et al., 2014). The fact that the *deltaC* stripe precedes the *her1* stripe only in the middle of the PSM is an additional indication for differential spatiotemporal modulation of time delays (Jülich et al, 2005).

Our results put the so-called desynchronization hypothesis, that the essential role of Notch signalling in somitogenesis is to maintain synchrony in neighbouring PSM oscillators (Jiang et al., 2000; Riedel-Kruse et al., 2007; Özbudak & Lewis, 2008; Liao & Oates, 2016), into perspective: Notch signalling also regulates the collective dynamics of cellular oscillators in terms of period and amplitude – and thereby shapes the travelling wave pattern.

Whether the oscillatory dynamics across the PSM are controlled by positional or temporal information is an open question (Oates et al., 2012; Shih et al., 2015). The ‘gradient by inheritance’ model suggests that *fgf8*, and possibly also *wnt3A*, are transcribed only in the tail bud and their mRNA is gradually decaying in the presomitic cells that are left behind by the proliferating tail bud (Dubrulle & Pourquié, 2004, Aulehla & Pourquié, 2010; Bajard et al., 2014). If one of these two signalling pathways would be involved in the shortening of the autorepression and coupling time delays, this would explain how temporal information of mRNA decay progression is translated into positional information by successive cell flow.

### 4.3. Unifying hypotheses for the role of time delays in Notch-mediated oscillation entrainment

A wide variety of time-delayed coupling phenomena related to the one investigated here, are currently studied across the natural sciences (Sadeghi & Valizadeh, 2014; Pavlides et al., 2015; Vanag et al., 2016; Wetzel et al., 2017). Recently, a unifying hypothesis for the time-delayed coupling mediated by Notch in neurogenesis and somitogenesis has been proposed (Shimojo et al., 2016; Shimojo & Kageyama, 2016): a difference in coupling delays between biological tissues could explain why dynamical salt-and-pepper patterning is observed in the developing brain and synchronization is observed in the PSM (Shimojo et al., 2016; Shimojo & Kageyama, 2016). This hypothesis has been based on the observation that mice mutants with different gene lengths, and therefore different transcriptional delays, exhibited oscillation death in the PSM on a tissue level (Shimojo & Kageyama, 2016). Oscillation death could theoretically occur between the regions of synchronization and anti-synchronization (Shimojo & Kageyama, 2016), but is not observed in our model of the zebrafish PSM oscillator (Figure 5B). Based on our model, we hypothesize that tissue-level oscillations are damped in coupling delay mutants by the shift to a dynamical salt-and-pepper patterning regime (Figure 4C and 4D; Figure 5). To gain a broader understanding, single cell oscillations remain to be tracked under these intriguing new experimental conditions with altered delay times (Shimojo & Kageyama, 2016). But importantly, the idea that the coupling delay varies the mode of entrainment in different tissues could apply in an even wider scope: it has been shown that during angiogenesis, upregulation of Dll4 by high Vegf signalling leads to a switch from dynamical salt-and-pepper patterning, required for tip versus stalk fate selection, to pathological synchronization of Dll4-Notch dynamics, leading to vessel expansion (Ubezio et al., 2016). Within our theoretical framework, such a switch would imply that Dll4 upregulation is modulating the relative length of the collective period and the coupling delay of the cellular oscillators (illustrated in Figure 5C), driving the system into a region of synchronization (Figure 6A). The sensitivity analysis of the emergent collective period (Figure 6B, Supplementary figure S2 and S3) suggests that time delays are the most potent regulators – and their modulation could be responsible for a switch in the mode of oscillation entrainment. Time delays might control both, the oscillatory dynamics within the PSM and the varying synchronization behaviour among different tissues. The molecular mechanisms that lead to these spatiotemporal gradients in time delays are still to be determined and should be the focus of future experimental studies.

Altogether, these recent advances the fields of somitogenesis, neurogenesis and angiogenesis support the view that time-delayed coupling of cellular oscillators via juxtacrine Notch signalling is a fundamental principle of developmental biology (Shimojo & Kageyama, 2016). The coupling behaviour controls whether cells differentiate collectively as in the PSM or whether a number of individual cells differentiate, as it is the case during the formation of the cerebral cortex or the branching of blood vessels. Additionally, the cellular oscillators in the PSM indirectly define the rhythmicity of segmentation via the formation of travelling waves.

### 4.4 Conclusions

We consider the *her7:hes6* double mutant as an ideal biological system to study somitogenesis mechanistically, because it carries a parsimonious PSM oscillator. Our model of this viable mutant characterizes how zebrafish PSM cells establish *her* oscillations, how they entrain their periods, synchronize locally, and how they adapt their behaviour collectively in the presence of spatiotemporal time delay gradients. In somitogenesis, a number of open questions still remain: How does the travelling wave pattern scale with the body and tissue size (Cooke, 1975; Tam, 1981; Schröter et al., 2008; Lauschke et al., 2013)? Is the travelling wave shaped by temporal or spatial cues (Oates et al., 2012; Shih et al., 2015)? And how does the travelling wave encode for segmentation (Oates et al., 2012; Lauschke et al., 2013; Shih et al., 2015)? We offer a solid mathematical framework that might be suitable to accommodate future experimental observations to address these questions. Furthermore, our framework offers a theoretical basis to comparatively study how emergent properties of time-delayed coupling via Notch signalling coordinate differentiation events across different embryonic tissues.

## Supplementary Information

### 1. Heuristic approach to build intuition for autorepression and coupling

Lewis approximated the autonomous *her1* period as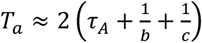. This implies that τ_A_< *T*_a_/*2* and consequently that autoinhibition must peak within the phase interval where the amplitude decreases. Consequently, for oscillatory curves that are symmetric in peaks and troughs, such as sinusoidal functions the autorepression delay τ_A_satisfies:

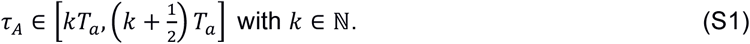

This condition is represented in Figure 3 with orange background colour.

Suppose there are two synchronized oscillators *A* and *B* with a collective period *T*_c_that mutually repress each other. They will remain synchronized only if the repression of *B* by *A* peaks within the oscillation phase of *B* where the amplitude is decreasing, and vice versa. Based on that, a relation between the common coupling delay τ_c_and the collective period *T*_c_analogous to Eq. S1 can be formulated – for oscillatory curves that are symmetric in peaks and troughs synchronized oscillators satisfy:

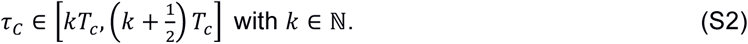

This condition is represented in Figure 5 with blue background colour.

**Supplementary figure S1:**
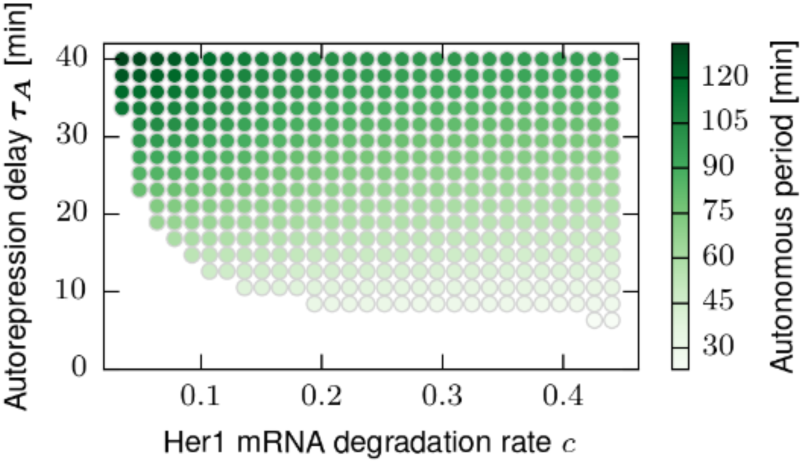
The autonomous period of Her1 is controlled by the autorepression delay and the degradation rates of the mRNA c (shown) and the protein b (equivalent to c). All data points correspond to 4×4 cell patches.

**Supplementary figure S2:**
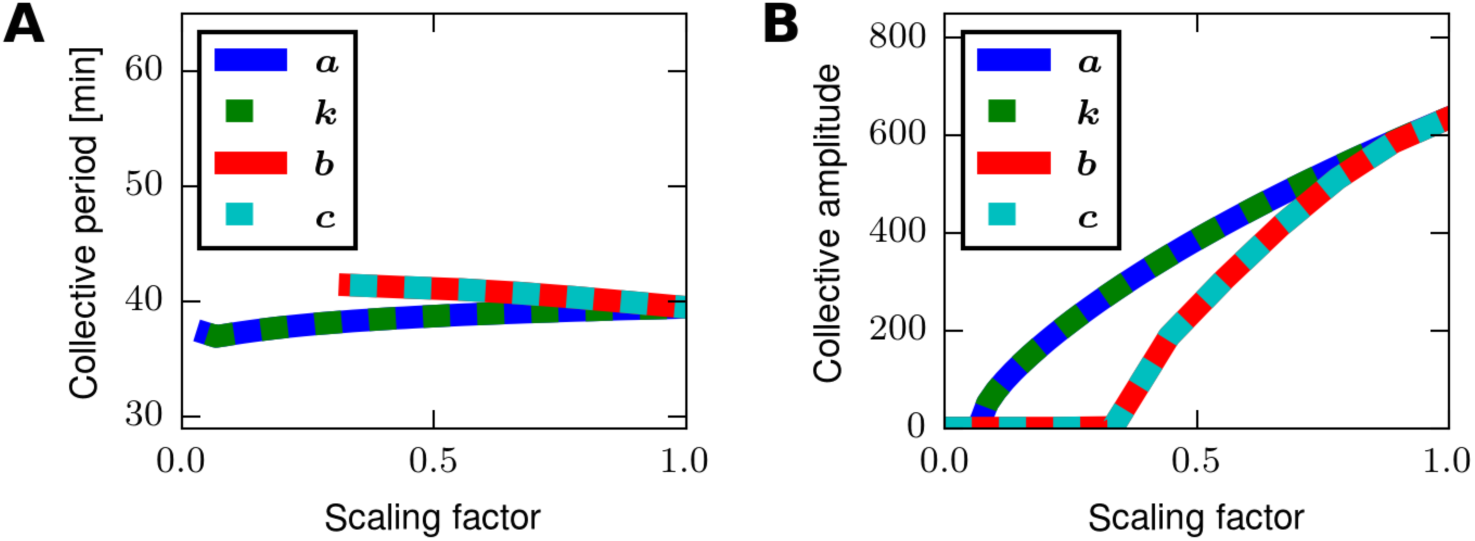
Sensitivity analysis of the kinetic rates associated with the Her1 oscillator in terms of collective period (A) and amplitude (B). Note, that the default values (scaling factor 1) are maximal estimates (Lewis, 2003). All data points correspond to 4×4 cell patches starting in a desynchronized initial condition (end point of Figure 4A).

**Supplementary figure S3:**
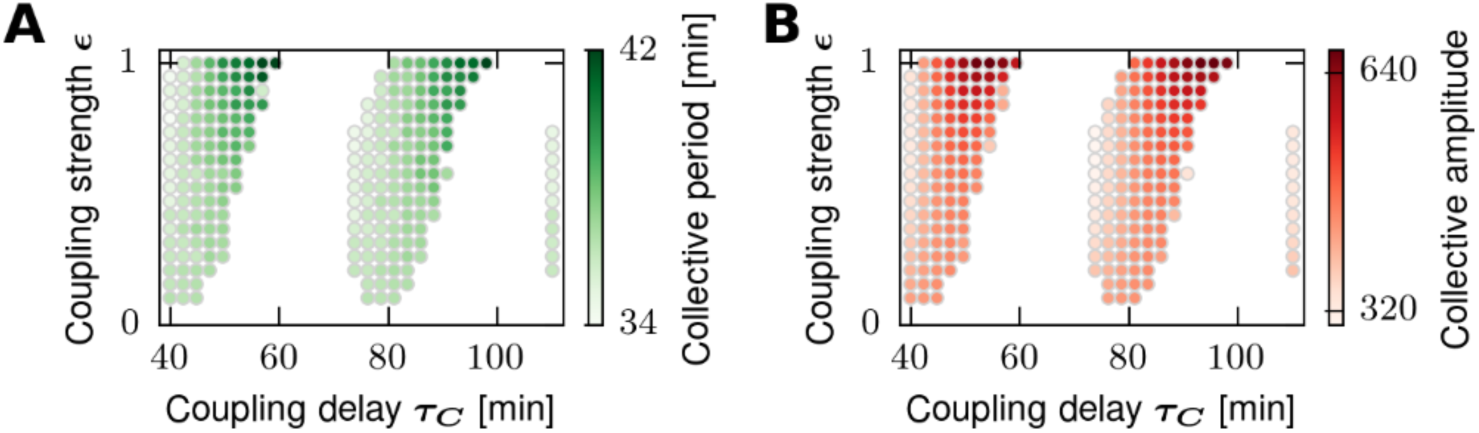
The role of the coupling strength. The collective periods (A) and amplitudes (B) of Her1 vary more with stronger coupling. The anti-synchronized states are hidden. The synchronized regions are moderately shifted with varying coupling strength. All data points correspond to 4×4 cell patches starting in a desynchronized initial condition (Figure 4A).

**Supplementary figure S4:**
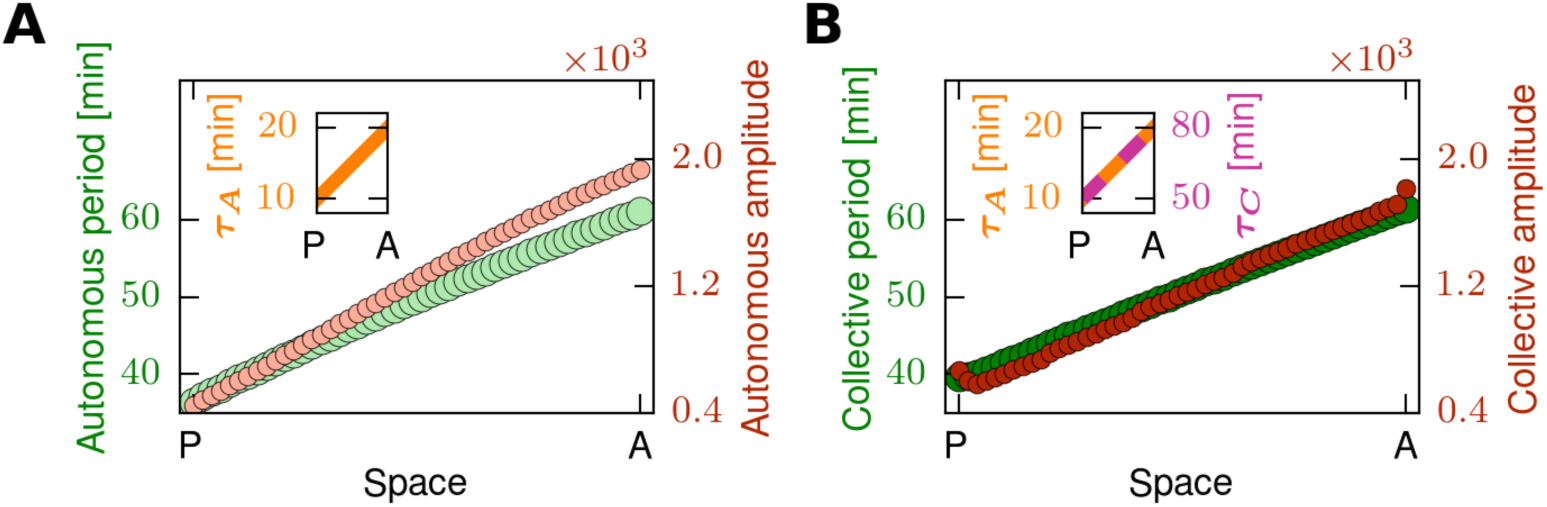
(A, B) Collective period and amplitude profiles of Her1 for static 4×50 cell tissues with default linear gradients in the autorepression delay τ_A_and the coupling delay τ_c_(embedded small plots) along the longer axis (posterior-anterior, P-A); recorded for autonomous (A) and coupled Her1 oscillators (B). In Figure 6, we have used 4×4 cell patches with homogeneous parameter values to approximate the PSM locally, similarly to the approach of others (Tiedemann et al., 2007; Ay et al., 2013). We have concluded that a spatiotemporal gradient in time delays (Figure 6B and 6C, black arrow) could explain the oscillatory dynamics of recorded across the PSM (Shih et al., 2015). Indeed, when imposing such a gradient spatially, we observe resultant linear gradients of collective period and amplitude of Her1, in space (A, B). This confirms that the local approximation with homogeneous 4×4 cell patches is reasonable. For the exemplary gradient, the period and amplitude pattern that we record in the autonomous (A) and the coupled scenario (B) are similar and roughly approximate the single cell wild type data measured in the PSM by others (Shih et al., 2015).

**Supplementary figure S5:**
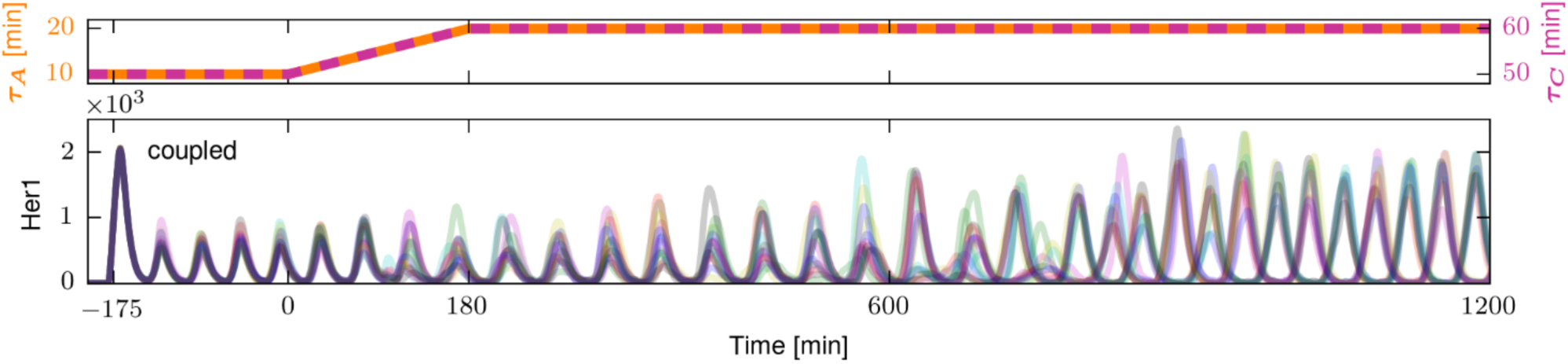
Simulation of coupling behaviour in presence of a temporal gradient in the autorepression delay τ_A_, which is included in the coupling delay τ_c_. Trajectories of coupled Her1 oscillations (bottom) in a group of 4×4 cells, in the presence of a temporal linear gradient in time delays between the timepoints 0 min and 180 min (top). The noise level corresponds to the value η = 10^−2^in Figure 4B.

## References

Aulehla, A., Wehrle, C., Brand-Saberi, B., Kemler, R., Gossler, A., Kanzler, B. and Herrmann, B. G.(2003). Wnt3a plays a major role in the segmentation clock controlling somitogenesis. Dev. Cell 4, 395–406.

Aulehla, A., Wiegraebe, W., Baubet, V., Wahl, M. B., Deng, C., Taketo, M., Lewandoski, M. and Pourquié, O.(2008). A β-catenin gradient links the clock and wavefront systems in mouse embryo segmentation. Nat. Cell Biol. 10, 186–193.

Aulehla, A., and Pourquié, O.(2010). Signaling gradients during paraxial mesoderm development. Cold Spring Harb. Perspect. Biol. 2, a000869.

Ay, A., Knierer, S., Sperlea, A., Holland, J. and Özbudak, E. M.(2013). Short-lived Her proteins drive robust synchronized oscillations in the zebrafish segmentation clock. Development 140, 3244–3253.

Ay, A., Holland, J., Sperlea, A., Devakanmalai, G. S., Knierer, S., Sangervasi, S., Stevenson, A. and Özbudak, E. M.(2014). Spatial gradients of protein-level time delays set the pace of the traveling segmentation clock waves. Development 141, 4158–4167.

Bajard, L., Morelli, L. G., Ares, S., Pécréaux, J., Jülicher, F. and Oates, A. C.(2014). Wnt-regulated dynamics of positional information in zebrafish somitogenesis. Development 141, 1381–1391.

Bessho, Y., Hirata, H., Masamizu, Y. and Kageyama, R.(2003). Periodic repression by the bHLH factor Hes7 is an essential mechanism for the somite segmentation clock. Genes Dev. 17, 1451–1456.

Clewley, R. H., Sherwood, W. E., LaMar, M. D. and Guckenheimer, J. M.(2007). PyDSTool, a software environment for dynamical systems modeling. URL http://pydstool.sourceforge.net

Cooke, J.(1975). Control of somite number during morphogenesis of a vertebrate, Xenopus laevis. Nature 254, 196–199.

Cooke, J.(1998). A gene that resuscitates a theory-somitogenesis and a molecular oscillator. Trends Genet. 14, 85–88.

Cooke, J. and Zeeman, E. C.(1976). A clock and wavefront model for control of the number of repeated structures during animal morphogenesis. J. Theor. Biol. 58, 455–476.

del Alamo, D., Rouault, H. and Schweisguth, F.(2011). Mechanism and significance of cis-inhibition in Notch signalling. Curr. Biol. 21, R40–47.

Delaune, E. A., François, P., Shih, N. P. and Amacher, S. L.(2012). Single-cell-resolution imaging of the impact of Notch signaling and mitosis on segmentation clock dynamics. Dev. Cell 23, 995–1005.

Dequeant, M. L., Glynn, E., Gaudenz, K., Wahl, M., Chen, J., Mushegian, A. and PourquiéO.(2006). A complex oscillating network of signaling genes underlies the mouse segmentation clock. Science 314, 1595–1598.

Diez del Corral, R., Olivera-Martinez, I., Goriely, A., Gale, E., Maden, M. and Storey, K.(2003). Opposing FGF and retinoid pathways control ventral neural pattern, neuronal differentiation, and segmentation during body axis extension. Neuron 40:65–79.

Dubrulle, J., McGrew, M. J. and Pourquié, O.(2001). FGF signaling controls somite boundary position and regulates segmentation clock control of spatiotemporal Hox gene activation. Cell 106, 219–232.

Dubrulle, J. and Pourquié, O.(2004). fgf8 mRNA decay establishes a gradient that couples axial elongation to patterning in the vertebrate embryo. Nature 427, 419–422.

Dunty, W.C.Jr., Biris, K. K., Chalamalasetty, R. B., Taketo, M. M., Lewandoski, M. and Yamaguchi, T. P.(2008). Wnt3a/β-catenin signaling controls posterior body development by coordinating mesoderm formation and segmentation. Development 135, 85–94.

Flunkert, V. and Schoell, E.(2009). Pydelay - a python tool for solving delay differential equations. arXiv:0911.1633.

Giudicelli, F., Özbudak, E. M., Wright, G. J. and Lewis, J.(2007). Setting the tempo in development: an investigation of the zebrafish somite clock mechanism. PLoS Biol. 5, e150.

Goldbeter, A., Gonze, D. and Pourquié, O.(2007). Sharp developmental thresholds defined through bistability by antagonistic gradients of retinoic acid and FGF signaling. Dev. Dyn. 236, 1495–1508.

Goldenfeld, N. and Kadanoff, L.(1999). Simple Lessons from Complexity. Science 284, 87.

Hanisch, A., Holder, M. V., Choorapoikayil, S., Gajewski, M., Özbudak, E. M. and LewisJ.(2013). The elongation rate of RNA polymerase II in zebrafish and its significance in the somite segmentation clock. Development 140, 444–453.

Harima, Y., Takashima, Y., Ueda, Y., Ohtsuka, T. and Kageyama, R.(2013) Accelerating the tempo of the segmentation clock by reducing the number of introns in the Hes7 gene. Cell Rep. 3, 1–7.

Henry, C. A., Urban, M. K., Dill, K. K., Merlie, J. P., Page, M. F., Kimmel, C. B. and Amacher, S. L.(2002). Two linked hairy/Enhancer of split-related zebrafish genes, her1 and her7, function together to refine alternating somite boundaries. Development 129, 3693–3704.

Herrgen, L., Ares, S., Morelli, L. G., Schröter, C., Jülicher, F. and Oates, A. C.(2010). Intercellular coupling regulates the period of the segmentation clock. Curr. Biol. 20, 1244–53.

Hester, S. D., Belmonte, J. M., Gens, J. S., Clendenon, S.G. and Glazier, J. A.(2011). A multi-cell, multi-scale model of vertebrate segmentation and somite formation. PLoS Comput Biol. 7, e1002155.

Hirata, H., Yoshiura, S., Ohtsuka, T., Bessho, Y., Harada, T., Yoshikawa, K. and KageyamaR.(2002). Oscillatory expression of the bHLH factor Hes1 regulated by a negative feedback loop. Science 298, 840–843.

Holley, S. A., Jülich, D., Rauch, G. J., Geisler, R. and Nüsslein-VolhardC.(2002). her1 and the notch pathway function within the oscillator mechanism that regulates zebrafish somitogenesis. Development 129, 1175–1183.

Hori, Y., Takada, M. and Hara, S.(2013). Biochemical oscillations in delayed negative cyclic feedback: Existence and profiles. Automatica 9, 2581–2591.

Horikawa, K., Ishimatsu, K., Yoshimoto, E., Kondo, S. and Takeda, H.(2006). Noise-resistant and synchronized oscillation of the segmentation clock. Nature 441, 719–723.

Hoyle, N. P. and Ish-Horowicz, D.(2013). Transcript processing and export kinetics are rate-limiting steps in expressing vertebrate segmentation clock genes. Proc. Natl. Acad. Sci. USA 110, E4316–E432.

Hubaud, A. and Pourquié, O.(2014). Signalling dynamics in vertebrate segmentation. Nat. Rev. Mol. Cell Biol. 15, 709–21.

Jaeger, J. and Goodwin, B. C.(2001). A cellular oscillator model for periodic pattern formation. J. Theor. Biol. 213, 171–181.

Jiang, Y.-J., Aerne, B. L., Smithers, L., Haddon, C., Ish-Horowicz, D. and Lewis, J.(2000). Notch signalling and the synchronization of the somite segmentation clock. Nature 408, 475–479.

Jenkins, R. P., Hanisch, A., Soza-Ried, C., Sahai, E. and Lewis, J.(2015). Stochastic Regulation of her1/7 Gene Expression Is the Source of Noise in the Zebrafish Somite Clock Counteracted by Notch Signalling. PLoS Comput. Biol., 11.

Jörg, D. J., Morelli, L. G., Soroldoni, D., Oates, A. C. and Jülicher, F.(2015). Continuum theory of gene expression waves during vertebrate segmentation. New J. Phys. 17, 093042.

Jörg, D. J., Oates, A. C. and Jülicher, F.(2016). Sequential pattern formation governed by signaling gradients. Phys. Biol. 13, 05LT03.

Jülich, D., Hwee Lim, C., Round, J., Nicolaije, C., Schroeder, J., Davies, A., Geisler, R., Lewis, J., Jiang, Y.-J. and Holley, S. A.(2005). beamter/deltaC and the role of Notch ligands in the zebrafish somite segmentation, hindbrain neurogenesis and hypochord differentiation. Dev. Biol. 286, 391–404.

Kærn, M., Menzinger, M. and Hunding, A.(2000). Segmentation and Somitogenesis derived from phase dynamics in growing oscillatory media. J. Theor. Biol. 207, 473–493.

Kageyama, R., Ohtsuka, T., Shimojo, H. and Imayoshi, I.(2008). Dynamic Notch signaling in neural progenitor cells and a revised view of lateral inhibition. Nature neuroscience 11, 1247–1251.

Kawamura, A., Koshida, S., Hijikata, H., Sakaguchi, T., Kondoh, H. and Takada, S.(2005). Zebrafish hairy/enhancer of split protein links FGF signaling to cyclic gene expression in the periodic segmentation of somites. Genes Dev. 19, 1156–1161.

Kim, J., Shin, D., Jung, S., Heslop-Harrison, P. and Cho, K.(2010). A design principle underlying the synchronization of oscillations in cellular systems.J. Cell Sci.123, 1537–1543.

Kim, W., Matsui, T., Yamao, M., Ishibashi, M., Tamada, K., Takumi, T., Kohno, K., Oba, S., Ishii, S., Sakumura, Y.et al.(2011). The period of the somite segmentation clock is sensitive to Notch activity. Mol. Biol. Cell 22, 3541–3549.

Krol, A., Roellig, D., Dequeant, M. L., Tassy, O., Glynn, E., Hattem, G., Mushegian, A., Oates, A. C. and Pourquié, O.(2011). Evolutionary plasticity of segmentation clock networks. Development 138, 2783–2792.

Lander, A. D.(2014). Making sense in biology: an appreciation of Julian Lewis. BMC Biol. 12, 57.

Lauschke, V. M., Tsiairis, C. D., François, P. and Aulehla, A.(2013). Scaling of embryonic patterning based on phase-gradient encoding. Nature 493, 101–105.

Lewis, J.(2003). Autoinhibition with transcriptional delay: a simple mechanism for the zebrafish somitogenesis. oscillator. Curr. Biol. 13, 1398–1408.

Liao, B. K., Jörg, D. J. and Oates, A. C.(2016). Faster embryonic segmentation through elevated Delta-Notch signalling. Nat. Commun.11861.

Liao, B. K. and Oates, A. C.(2016). Delta-Notch signalling in segmentation. Arthropod. Struct. Dev.[Epub ahead of print]. doi: 10.1016/j.asd.2016.11.007.

Mara, A., Schroeder, J., Chalouni, C. and Holley, S. A.(2007). Priming, initiation and synchronization of the segmentation clock by deltaD and deltaC.Nat. Cell Biol. 9, 523–530.

Maroto, M., Dale, J. K., Dequéant, M. L., Petit, A. C. and PourquiéO.(2005). Synchronised cycling gene oscillations in presomitic mesoderm cells require cell-cell contact. Int. J. Dev. Biol. 49, 309–315.

Masamizu, Y., Ohtsuka, T., Takashima, Y., Nagahara, H., Takenaka, Y., Yoshikawa, K., Okamura, H. and Kageyama, R.(2006). Real-time imaging of the somite segmentation clock: revelation of unstable oscillators in the individual presomitic mesoderm cells. Proc. Natl. Acad. Sci. USA 103, 1313–1318.

Matsuda, M. and ChitnisA. B.(2009). Interaction with Notch determines endocytosis of specific Delta ligands in zebrafish neural tissue. Development 136, 197–206.

Momiji, H. and Monk, N. A.(2009). Oscillatory Notch-pathway activity in a delay model of neuronal differentiation. Phys. Rev. E Stat. Nonlin. Soft Matter Phys. 80, 021930.

Monk, N. A.(2003). Oscillatory expression of Hes1, p53, and NF-κB driven by transcriptional time delays. Curr. Biol. 13, 1409–13.

Morelli, L. G., Ares, S., Herrgen, L., Schröter, C., Jülicher, F. and OatesA. C.(2009). Delayed coupling theory of vertebrate segmentation. HFSP J. 3, 55–66.

Murray, P.J., Maini, P. K., Baker, R.E.(2013). Modelling Delta-Notch perturbations during zebrafish somitogenesis. Dev. Biol.,373, 407–421

Naiche, L. A., Holder, N. and Lewandoski, M.(2011). FGF4 and FGF8 comprise the wavefront activity that controls somitogenesis.Proc. Natl. Acad. Sci. USA 108, 4018–4023.

Niebur, E., Schuster, H. G. and Kammen, D. M.(1991). Collective frequencies and metastability in networks of limit-cycle oscillators with time delay. Phys. Rev. Lett. 67, 2753–2756.

Niederreither, K., McCaffery, P., Dräger, U. C., Chambon, P. and Dollé, P.(1997). Restricted expression and retinoic acid-induced downregulation of the retinaldehyde dehydrogenase type 2 (RALDH-2) gene during mouse development. Mech. Dev. 62, 67–78.

Novak, B. and Tyson, J. J.(2008). Design principles of biochemical oscillators. Nat. Rev. Mol. Cell Biol. 9, 981–991.

Palmeirim, I., Henrique, D., Ish-Horowicz, D. and Pourquié, O.(1997). Avian hairy gene expression identifies a molecular clock linked to vertebrate segmentation and somitogenesis. Cell 91, 639–648.

Pavlides, A., Hogan, S. J. and Bogacz, R.(2015). Computational Models Describing Possible Mechanisms for Generation of Excessive Beta Oscillations in Parkinson’s Disease. PLoS Comput. Biol. 11, e1004609.

Oates, A. C. and Ho, R. K.(2002). Hairy/E(spl)-related (Her) genes are central components of the segmentation oscillator and display redundancy with the Delta/Notch signaling pathway in the formation of anterior segmental boundaries in the zebrafish. Development 129, 2929–2946.

Oates, A. C., Morelli, L. G. and Ares, S.(2012). Patterning embryos with oscillations: structure, function and dynamics of the vertebrate segmentation clock. Development 139, 625–39.

Oginuma, M., Niwa, Y., Chapman, D. L. and Saga, Y.(2008). Mesp2 and Tbx6 cooperatively create periodic patterns coupled with the clock machinery during mouse somitogenesis. Development 135, 2555–2562.

Özbudak, E. M. and Lewis, J.(2008). Notch signalling synchronizes the zebrafish segmentation clock but is not needed to create somite boundaries. PLoS Genet. 4, e15.

Riedel-KruseI. H., Müller, C. and OatesA. C.(2007). Synchrony dynamics during initiation, failure, and rescue of the segmentation clock. Science 317, 1911–5.

Sadeghi, S. and Valizadeh, A.(2014). Synchronization of delayed coupled neurons in presence of inhomogeneity. Journal of computational neuroscience 36, 55–66.

Saga, Y.(2012). The mechanism of somite formation in mice. Curr. Opin. Genet. Dev. 22, 331–8.

Saga, Y., Hata, N., Koseki, H. and Taketo, M. M.(1997). Mesp2: a novel mouse gene expressed in the presegmented mesoderm and essential for segmentation initiation. Genes Dev. 11, 1827–1839.

Sawada, A., Fritz, A., Jiang, Y.-J., Yamamoto, A., Yamasu, K., Kuroiwa, A., Saga, Y. and Takeda, H.(2000). Zebrafish Mesp family genes, mesp-a and mesp-b are segmentally expressed in the presomitic mesoderm, and Mesp-b confers the anterior identity to the developing somites. Development 127, 1691–1702.

Sawada, A., Shinya, M., Jiang, Y.-J., Kawakami, A., Kuroiwa, A. and Takeda, H.(2001). Fgf/MAPK signalling is a crucial positional cue in somite boundary formation.Development 128, 4873–4880.

Schröter, C., Herrgen, L., Cardona, A., Brouhard, G. J., Feldman, B. and OatesA. C.(2008). Dynamics of zebrafish somitogenesis. Dev. Dyn. 237, 545–553.

Schröter, C., Ares, S., Morelli, L. G., Isakova, A., Hens, K., Soroldoni, D., Gajewski, M., Jülicher, F., Maerkl, S. J., Deplancke, B.et al.(2012). Topology and dynamics of the zebrafish segmentation clock core circuit. PLoS Biol. 10, e1001364.

Shih, N. P., François, P., Delaune, E. A. and Amacher, S. L.(2015). Dynamics of the slowing segmentation clock reveal alternating two-segment periodicity.Development 142, 1785–1793.

Shimojo, H., Isomura, A., Ohtsuka, T., Kori, H., Miyachi, H. and R. Kageyama. (2016). Oscillatory control of Delta-like1 in cell interactions regulate dynamic gene expression and tissue morphogenesis. Genes Dev. 30, 102–116.

Shimojo, H. and Kageyama, R.(2016). Oscillatory control of Delta-like1 in somitogenesis and neurogenesis: a unified model for different oscillatory dynamics. Semin. Cell Dev. Biol. 49, 76–82.

Shimojo, H., Ohtsuka, T. and Kageyama, R.(2008). Oscillations in notch signaling regulate maintenance of neural progenitors.Neuron 58, 52–64.

Soroldoni, D., Jörg, D. J., Morelli, L. G., Richmond, D. L., Schindelin, J., Jülicher, F. and Oates, A. C.(2014). A Doppler effect in embryonic pattern formation. Science 345, 222–5.

Soza-Ried, C., Öztürk, E., Ish-Horowicz, D. and Lewis, J.(2014). Pulses of Notch activation synchronise oscillating somite cells and entrain the zebrafish segmentation clock. Development 141, 1780–1788.

Takahashi, Y., Koizumi, K., Takagi, A., Kitajima, S., Inoue, T., Koseki, H. and Saga, Y.(2000). Mesp2 initiates somite segmentation through the Notch signalling pathway. Nature Genet. 25, 390–396.

Takashima, Y., Ohtsuka, T., González, A., Miyachi, H. and Kageyama, R.(2011). Intronic delay is essential for oscillatory expression in the segmentation clock. Proc. Natl. Acad. Sci. USA 108, 3300–3305.

Tam, P. P.(1981). The control of somitogenesis in mouse embryos. J. Embryol. Exp. Morphol. 65, 103–128.

Tiedemann, H. B., Schneltzer, E., Zeiser, S., Rubio-Aliaga, I., Wurst, W., Beckers, J., Przemeck, G. K. and Hrabé de Angelis, M.(2007). Cell-based simulation of dynamic expression patterns in the presomitic mesoderm. J. Theor. Biol. 248, 120–129.

Tiedemann, H. B., Schneltzer, E., Zeiser, S., Hoesel, B., Beckers, J., Przemeck, G. K. and Hrabé de Angelis, M.(2012) From dynamic expression patterns to boundary formation in the presomitic mesoderm. PLoS Comput. Biol. 8, e1002586.

Rost, F., Eugster, C., Schröter, C., Oates, A. C. and Brusch, L.(2014). Chevron formation of the zebrafish muscle segments. J. Exp. Biol. 217, 3870–3882.

Ubezio, B., Blanco, R. A., Geudens, I., Stanchi, F., Mathivet, T., Jones, M. L., Ragab, A., Bentley, K. and GerhardtH.(2016). Synchronization of endothelial Dll4-Notch dynamics switch blood vessels from branching to expansion. eLife 5, e12167.

Uriu, K., Morishita, Y. and Iwasa, Y.(2009). Traveling wave formation in vertebrate segmentation. J. Theor. Biol. 257, 385–396.

Vanag, V. K., Smelov, P. S. and Klinshov, V. V.(2016). Dynamical regimes of four almost identical chemical oscillators coupled via pulse inhibitory coupling with time delay. Phys. Chem. Chem. Phys. 18, 5509–20.

Wang, Y. Q., Hori, Y., Hara, S. and Doyle, F. J.(2014). Intercellular delay regulates the collective period of repressively coupled gene regulatory oscillator networks. IEEE Trans. Autom. Control 59, 211–216.

Wang, Y. Q., Hori, Y., Hara, S. and Doyle, F. J.(2015). Collective oscillation period of inter-coupled biological negative cyclic feedback oscillators. IEEE Trans. Autom. Control 60, 1392–1397.

Wanglar, C., Takahashi, J., Yabe, T. and Takada, S.(2014). Tbx protein level critical for clock-mediated somite positioning is regulated through interaction between Tbx and Ripply. PLoS ONE 9, e107928.

Ward, A. B. and Mehta, R. S.(2011). Axial elongation in fishes: using morphological approaches to elucidate developmental mechanisms in studying body shape. Integr. Comp. Biol. 50, 1106–1119.

Webb, A. B., Lengyel, I. M., Jörg, D. J., Valentin, G., Jülicher, F., Morelli, L. G., and Oates, A. C.(2016). Persistence, period and precision of autonomous cellular oscillators from the zebrafish segmentation clock. eLife 5, e08438.

Wetzel, L., Jörg, D. J., Pollakis, A., Rave, W., Fettweis, G. and Jülicher, F.(2017). Self-organized synchronization of digital phase-locked loops with delayed coupling in theory and experiment. PLoS ONE 12, e0171590.

Wright, G. J., Giudicelli, F., Soza-Ried, C., Hanisch, A., Ariza-McNaughton, L. and Lewis, J.(2011). DeltaC and DeltaD interact as Notch ligands in the zebrafish segmentation clock. Development 138, 2947–2956.

Yabe, T. and Takada, S.(2016). Molecular mechanism for cyclic generation of somites: lessons from mice and zebrafish. Dev. Growth Differ. 58, 31–42.

Yasuhiko, Y., Haraguchi, S., Kitajima, S., Takahashi, Y., Kanno, J. and Saga, Y.(2006). Tbx6-mediated Notch signaling controls somite-specific Mesp2 expression. Proc. Natl. Acad. Sci. USA 103, 3651–3656.

